# *Leishmania amazonensis* amastigotes invade non-phagocytic cells via highly localized parasite-induced actin remodeling

**DOI:** 10.1101/2025.10.13.681999

**Authors:** Thamires Queiroz-Oliveira, Laura Valéria Rios-Barros, Anna Luiza Silva-Moreira, Isadora Vieira Santos-Brasil, Julyanna Oliveira-Castro, Juliana Almeida, Marcos André Vannier-Santos, Jane Lima-Santos, Maria Fátima Horta, Thiago Castro-Gomes

**Author notes:** (Thamires Queiroz-Oliveira); (Laura Valéria Rios-Barros); (Anna Luiza Moreira-Silva); (Isadora Vieira Santos-Brasil); (Julyanna Oliveira-Castro); (Juliana Almeida); (Marcos André Vannier-Santos); (Jane Lima-Santos); (Maria Fátima Horta); (Thiago Castro-Gomes). Corresponding author: Thiago de Castro Gomes, Ph.D, Departamento de Parasitologia, Bloco L4 Sala 244 Instituto de Ciências Biológicas (ICB) Universidade Federal de Minas Gerais (UFMG) Av. Antônio Carlos, 6627 – Campus Pampulha Belo Horizonte, MG 31270-901, Brazil.

## Abstract

Intracellular parasites are pathogens that must invade and persist within host cells. In the case of *Leishmania* spp., it is generally assumed that the parasite must be phagocytosed to further establish residence within professional phagocytic cells. However, several studies report the presence of *Leishmania* spp. in non-phagocytic cells, highlighting their ability to enter cells independently of classical phagocytosis. Indeed, we have recently demonstrated that promastigotes, the infective form transmitted to hosts during the bite of the insect vector, subvert a lysosome-exocytosis-dependent membrane repair pathway to invade fibroblasts. Here, we investigate the invasion of non-phagocytic cells by *L. amazonensis* amastigotes, the infective form directly responsible for host-to-vector transmission, disease generation and parasite dissemination during infection in humans and other mammalian hosts. Our results show that amastigotes rapidly induce their entry into cells lacking classical phagocytic capability, where they survive, multiply, and persist. Invasion depends on parasite-induced actin remodeling confined to the parasite-host plasma membrane contact site, with localized recruitment of Rho GTPases that fuel actin polymerization at invasion foci. Our findings highlight the remarkable ability of *Leishmania* amastigotes to induce their own internalization into virtually any cell type, bypassing the need for conventional phagocytosis. This property may profoundly influence parasite biology by enabling amastigotes to cross cellular barriers, disseminate, and silently establish infection in diverse host cells. Importantly, when considering non-phagocytic cells, our results demonstrate that *Leishmania amazonensis* employs distinct, stage-specific invasion mechanisms: promastigotes co-opt host cell lysosomes, whereas amastigotes depend on F-actin dynamics.

**SUMMARY:** Here we elucidate key aspects of the mechanism by which *Leishmania amazonensis* amastigotes invade non-phagocytic cells.

## INTRODUCTION

Parasites of the genus *Leishmania* comprise several protozoan species that cause a group of human diseases collectively known as leishmaniases. These infections are considered the second deadliest parasitic diseases, after malaria (1). Leishmaniases are endemic in many countries across Latin America, Europe, Africa, the Middle East, and South Asia, putting more than 1 billion people at risk, with more than 12 million currently infected (2) and around 1 million new cases each year (3). The outcome of leishmaniases depends on the parasite species and strain characteristics and the host’s immunological response to infection. The complexity of the host-parasite interaction underlies the broad spectrum of clinical manifestations associated with the disease. Some *Leishmania* species, such as *L. braziliensis*, *L. major* and *L. amazonensis*, primarily cause cutaneous leishmaniasis. Other species, most notably *L. infantum* and *L. donovani*, cause visceral leishmaniasis, the most severe form of the disease, which affects internal organs such as spleen, liver and bone marrow.

Parasites of the genus *Leishmania* alternate between two main evolutionary forms during their life cycle. Promastigotes, the flagellated extracellular form, reside and reproduce in the midgut of hematophagous female sandflies, while amastigotes, the round-shaped intracellular form, inhabit vacuoles within mammalian host cells. During a blood meal, infected sandflies inoculate promastigotes into the host’s dermis, where they are subsequently engulfed by neutrophils and ultimately by macrophages (4). Inside host cells, parasites transform into amastigotes and replicate intracellularly. Amastigotes are the main drivers of leishmaniasis pathogenesis as they multiply and invade new cells via poorly understood mechanisms (5), thereby amplifying infection. Because amastigotes are primarily found within macrophages in host lesions, these cells have been the most extensively studied and considered the standard model of in vitro infections. However, *Leishmania* spp. amastigotes have also been observed inside non-phagocytic cells including fibroblasts, epithelial cells, muscle cells and others, both in natural and experimental infections (6–23). Thus, contrary to the prevailing view that infection requires phagocytosis, *Leishmania* amastigotes can invade non-phagocytic cells.

Despite growing evidence that *Leishmania* spp. infect a variety of cells beyond professional phagocytes, the mechanisms involved in the process of cell invasion as well as the role of non-phagocytic cells in leishmaniasis has long been overlooked. We recently identified the mechanistic hallmarks of promastigote entry into such cells, demonstrating its dependence on the recruitment of host cell lysosomes involved in plasma membrane repair (24), a mechanism also exploited by *Trypanosoma cruzi* during invasion (25). In contrast, the mechanisms enabling amastigote invasion of non-phagocytic cells remain largely unknown. This underexplored capacity may profoundly influence pathogenesis by establishing protected niches that foster immune evasion, drug resistance, and disease relapse.

In this study, we investigated the mechanisms enabling *L. amazonensis* amastigotes to invade non-phagocytic cells. Our findings show that amastigotes induce localized actin polymerization, leading to intense plasma membrane ruffling, with RhoA and Rac1 recruited to sites of entry. Remarkably, this mechanism is entirely distinct from that of promastigotes, underscoring that *L. amazonensis* employs stage-specific invasion strategies that may critically shape infection establishment, progression and host-to-vector infection.

## MATERIAL AND METHODS

### 1 – Parasites and host cells

#### 1.1 – Cultivation of axenic amastigotes and promastigotes of *L. amazonensis*

The PH8 strain of *Leishmania (Leishmania) amazonensis* (*LLa*) used in this study was kindly provided by Dr. Maria Norma Melo (Departamento de Parasitologia, Universidade Federal de Minas Gerais, Belo Horizonte, Brazil). *L. amazonensis* expressing Red Fluorescent Protein (*LLa*-RFP) was provided by Dr. David Sacks (NIH, Bethesda, USA) and cultured as previously described by Carneiro et al. (2018) (26). Parasites were maintained at 24°C in M199 medium (GIBCO) supplemented with 10% heat-inactivated fetal bovine serum (FBS) (GIBCO), 100 U/ml penicillin, and 100 μg/ml streptomycin (GIBCO). *LLa*-RFP promastigotes were cultured under the same conditions as wild-type (WT) promastigotes, with the addition of 50 μg/ml geneticin (G418, Life Technologies) for the selection of RFP-expressing parasites. Cultures were initiated with 1 x 10^6^ parasites/ml and grown for 4-6 days, resulting in an enrichment of infective metacyclic promastigotes which were purified using a Ficoll gradient, as described by Späth and Beverley (2001) (27).

To obtain axenic *LLa*-RFP amastigotes, we followed the protocol described by Sarkar et al. (2018)(28), which employs M199 medium (identical to that used for promastigotes) supplemented with 0.25% glucose, 0.5% trypticase (peptone), 40 mM succinic acid, 1% adenine (10 mM), 1% L-glutamine, and 2 mM hemin. Starting with fifth-day promastigote cultures, the parasites were transferred into the amastigote medium and cultured at 34°C. After transformation into amastigotes, parasites were maintained using the same inoculum and culture conditions as those used for promastigotes, but at 34°C. Healthy axenic amastigotes appeared oval in shape, lacked visible flagella, and had a distinct plasma membrane, well-defined nucleus, and kinetoplasts, as confirmed by Leishman staining. The transformation of the parasites from promastigotes to amastigotes was monitored using microscopy and viability analysis (Fig. S1). Additionally, their ability to shift between these two evolutive forms in vitro and their capacity to infect cells were evaluated (Fig. S1 and S2).

#### 1.2 – Isolation of intracellular amastigotes of *L. amazonensis*

For the assays with non-axenic amastigotes, macrophage-like RAW cells were cultured at 37°C, in a 5% CO₂ atmosphere, in 9 mm² culture plates (KASVI) at a density of 4 x 10⁵ cells per plate, seeded 24 hours prior to infection. To infect RAW cells, metacyclic *LLa*-RFP promastigotes were added at a ratio of 25 parasites per macrophage and incubated for 48 hours at 37°C, 5% CO₂. After this period, cells contained amastigote-filled vacuoles. To isolate and obtain intracellular amastigotes, infected RAW cells were washed three times with PBS, then incubated for 30 seconds with lysis solution (DMEM containing 0.05% SDS), followed by the addition of DMEM to dilute the SDS, as described by Jain et al. (2012)(29). The non-axenic amastigotes were washed in PBS and used in the indicated experiments.

#### 1.3 – Cultivation of mammalian host cells

Mouse embryonic fibroblasts (MEFs), Human Umbilical Vein Endothelial Cells (HUVEC), Henrietta Lacks cells (HeLa), Human Hepatocellular Carcinoma cell line G2 (HepG2), or RAW cells were cultured in DMEM (Gibco) with 10% heat-inactivated FBS (Gibco) at 37°C in a 5% CO₂ atmosphere. Cultures were seeded every 48 hours and plated 24 h before experiments on culture dishes (Sarstedt) or glass coverslips, as needed. Cells were detached using 0.5% trypsin for MEFs, HUVEC, HeLa, HepG2 and cell scrapers for RAW cells. Sub-confluent cultures were used for infection experiments, followed by analysis via fluorescence microscopy or flow cytometry. In these experiments, cells were plated in 6-well dishes (Kasvi) at 3 × 10⁵ cells per well, with round coverslips placed in wells before plating for immunofluorescence. All cell lines were routinely checked for contamination.

### 2 – Viability tests

To assess the viability of cultivated parasites, both in amastigote and promastigote forms, two different tests were employed: erythrosine staining and the metabolic MTT assay (Fig. S1D-E). The erythrosine technique only stains dead parasites, allowing for counting of viable parasites in a Neubauer chamber. The percentage of viable parasites was then calculated. The MTT test, a colorimetric assay, relies on the ability of viable parasites to reduce tetrazolium salt to an insoluble formazan product, which is quantified using a spectrophotometer. This protocol, commonly used for mammalian cells, was adapted for *Leishmania* as described by Dutta et al. (2005) (30). Only cultures with over 90% viability were used in infection experiments (Fig. S2).

### 3 – Infection experiments

*LLa* amastigotes (axenic or intracellular) or metacyclic promastigotes were used in all experiments. Parasites were added to dish-adherent MEFs in DMEM containing 10% heat-inactivated FBS (GIBCO), and the plates were centrifuged at 500 x g for 10 minutes at 15°C to synchronize parasite contact with the cell monolayers. The cells were then incubated at 37°C in a 5% CO2 atmosphere for the indicated periods. All experiments were conducted with a multiplicity of infection of 25 parasites per cell.

In some assays, the parasites were fixed in 4% paraformaldehyde (PFA) for 15 minutes to determine whether cells were capable of actively capturing dead parasites, or if parasite viability was critical for cell invasion. For other assays, drugs such as Cytochalasin-D (SIGMA) and ROCK inhibitor (SIGMA) were used to assess the role of the F-actin cytoskeleton and its polymerization in the invasion process. Cells were pretreated with 10 µM of these drugs for 30 and 120 minutes, respectively, before being added to the culture medium. After treatment, the culture medium was replaced, and cells were incubated with the parasites for the specified duration. The cells were then detached using 0.25% trypsin, which also detaches parasites bound to the cells but not internalized. Infection was quantified using flow cytometry (FACS). For microscopy-based assays, cells grown on coverslips were fixed with 4% PFA for 15 minutes, followed by immunostaining with appropriate probes. In some cases, microscopy data were quantified by manual counting after immunostaining with anti-LAMP1 antibodies, which enabled the counting of parasites inside parasitophorous vacuoles.

### 4 – Cell staining

**Leishman staining** – Slides containing adhered promastigotes were fixed with methanol for 1 minute. Subsequently, Leishman stain solution (Newprov), diluted in buffered water at a 1:2 ratio, was applied to the slide and left to react for 20 minutes. After staining, the slides were carefully rinsed with buffered water and air-dried at room temperature. **Giemsa staining** – The slides were previously fixed with methanol for 2 minutes. After drying, a 10% Giemsa solution (Newprov) (diluted in buffered water) was applied and allowed to react for 15 minutes. Next, the slides were washed with water and left to dry. **Panoptic staining** – A panoptic staining kit (Laborclin) was used in some experiments. This kit consists of a hematological differential staining protocol performed on dead cells, based on the May-Grünwald-Giemsa method. Slides were first immersed for 30 seconds in the fixative solution (stain 1), followed by 2 minutes in the eosin solution (stain 2), and finally for 5 seconds in the azur stain solution (stain 3). After staining, the slides were rinsed with distilled water, air-dried at room temperature, mounted with Entellan (Merck) and subsequently analyzed under a light microscope.

### 5 – Cell labeling for fluorescence microscopy

**Immunofluorescence and fluorescent probes** – Sub-confluent cell monolayers were infected with LLa-RFP amastigotes or promastigotes for the indicated time periods and fixed with 4% paraformaldehyde. Preparations were then blocked and permeabilized using PBS containing 2% BSA and 0.5% saponin. The following antibodies or compounds were used for cell labelling: rat anti-LAMP-1 IgG (1D4B) (obtained from the Developmental Studies Hybridoma Bank) to label lysosomes (1:50, diluted in the same buffer), rabbit anti-RhoA (Thermo) to label RhoA IgG (1:300, diluted in the same buffer), mouse Anti-Rac1 IgG (BD – Transduction Laboratories) to label Rac1 (1:100, diluted in the same buffer), and mouse anti-CD45 FITC (BD Pharmingen) to label CD45 in macrophages-like cells (RAW) (1:50 diluted in the same buffer). For F-actin labeling, Alexa-488 or Alexa-633-conjugated phalloidin (Life Technologies) was used following the manufacturer’s protocol, with a 1:250 dilution (6 µM stock solution) in 0.1% Triton. The final concentration of phalloidin used for F-actin staining was 150 nM. After washing, where appropriate, preparations were incubated for 30 minutes with 1:250 Alexa-Fluor-conjugated secondary antibodies (Life Technologies). All preparations were stained with DAPI to visualize nuclei. Coverslips were mounted on microscope slides using Prolong-Glass anti-fade reagent (Life Technologies) and analyzed by fluorescence microscopy. Images were acquired and analyzed using Q-Capture software or Zen Software (ZEISS), depending on the experiment, as indicated.

### 6 – Fluorescence microscopy, image acquisition, treatment and quantitative analysis

Zeiss ApoTome, LSM 880 confocal, and Olympus BX-60 upright fluorescence microscopes were used throughout the experiments, as indicated in each specific figure legend. Z-stack images were acquired using a Zeiss LSM 880 confocal microscope and Apotome microscopes. The Z-stack range was defined manually by selecting the upper and lower focal planes of the sample, adjusted to the optimal slide thickness. Optical slices were collected at 0.5 μm intervals using either a 40x or 63x oil immersion objective, with the pinhole set to 1 Airy unit and an optimal pixel size. All imaging parameters (laser intensity, gain, and detector settings) were kept constant throughout acquisition. Image processing and projections were performed using ZEN Blue software (ZEISS). 2.5D topographical representation was generated using ZEN Blue (ZEISS). Intensity-based surface plots were constructed from single-plane confocal images. This visualization allowed for assessment of signal distribution and surface topology of the labeled structures. Both Zen Blue (ZEISS) and Image-J were used to treat images and perform quantifications. To quantify F-actin or LAMP-1 accumulation in parasitophorous vacuoles, we used the methodology previously described by our group to measure lysosome spreading within mammalian cells (31). For amastigotes, the parasite radius was used to define the vacuolar area surrounding each parasite, where the intensity of F-actin or LAMP-1 was then measured.

### 7 – Spectral Imaging with CHS1 Spectral Detector

Spectral imaging was performed using a Zeiss LSM 880 confocal microscope equipped with the Channel Spectral Detector 1 (CHS1). Samples were prepared on coverslips and mounted using a Prolong-Glass antifade reagent. Imaging was conducted using a 63x oil immersion objective. Fluorophores were excited sequentially using five laser lines: 405 nm, 488 nm, 546 nm, 583 nm, and 633 nm. Emission spectra were acquired in lambda mode over a defined spectral range appropriate for each fluorophore. Spectral detection was performed via the CHS1 detector, which allows simultaneous acquisition of up to 32 spectral channels per pixel. To ensure consistency, the pinhole size was set to 1 Airy unit, and acquisition settings (laser power, detector gain, offset, averaging, and scan speed) were kept constant across all imaging sessions. Linear spectral unmixing was applied using the ZEN Blue software (ZEISS), based on reference spectra acquired from single-labeled controls. All images were acquired in the same session to minimize variability due to laser fluctuations or alignment shifts (Fig. 7C-G and Fig. S9).

### 8 – Scanning Electron Microscopy (SEM)

After the infection assay, host cells and parasites were fixed in 2.5% glutaraldehyde, 2% formaldehyde, and 2.5 mM CaCl₂ in 0.1 M sodium cacodylate buffer (pH 7.2). The samples were then post-fixed in 1% OsO₄, 0.8% potassium ferricyanide, and 2.5 mM CaCl₂ in the same buffer for 60 minutes. Following this, the samples were dehydrated through a series of ethanol solutions, subjected to critical point drying using a Balzers apparatus, and mounted on bases. The samples were then observed under a JEOL 5310 scanning electron microscope.

### 9 – Quantification and visualization of infection

**FACS:** To quantify the infection rate, we used the *LLa*-RFP strain described above. After infection experiments, cells were washed and treated with 0.25% trypsin (Gibco) to detach host cells and non-internalized parasites. The cell population was then immediately analyzed by flow cytometry using a FACSCAN II (Becton Dickinson). A total of 10,000 events (MEFs) were analyzed, and data were processed using FlowJo software. The gating strategy used is demonstrated in the supplementary material (Fig. S3). To compare infection rates under different treatments, we used the mean fluorescence intensity (MFI) obtained from FACS analysis or manual counting of internalized parasites under the microscope, as indicated. **Light Microscopy:** Infected cells were visualized using a BX60 Upright Compound Fluorescence Microscope (Olympus) after staining with a hematoxylin-eosin panoptic kit (Laborclin) or GIEMSA. Cells were mounted on microscopy slides using Entellan (Merk), and images were obtained with Q-CapturePro Software. **Fluorescence Microscopy:** Cells labeled with fluorophore-conjugated antibodies or probes were analyzed with a BX60 Upright Compound Fluorescence Microscope (Olympus), Axio Imager ApoTome2 Microscope (Zeiss), Axio Imager ApoTome3 Microscope (Zeiss), and LSM 880 Confocal Microscope (Zeiss) to obtain confocal images (Z-stack, 2.5D image and Spectral Imaging (Detector CHS1)).

### 10 – Repeats, statistics, and extra images

Each experiment was performed in simple duplicates and repeated in at least three independent biological replicates, with representative results shown. Statistical analyses, including p-value calculations, were performed as indicated in the figure legends, where applicable. Additional experimental images and extra controls are available in the supplementary material.

## RESULTS

### *L. amazonensis* amastigotes invade MEFs

To assess the ability of amastigote forms to invade MEFs, we incubated them with RFP-expressing axenic amastigotes of *L. amazonensis* (*LLa*-AMA) and quantified infection over time using FACS (Fig. 1A), following the gating strategy shown in the supplementary material (Fig. S3). The results revealed a progressive increase in parasite invasion, from the initial interaction with the cell monolayer to later time points, culminating in an infection rate of 74% by the end of the experiment, 4 h after parasite inoculation. Fluorescence microscopy analysis with F-actin labeling, performed 4 h after infection to capture later stages, revealed the presence of fully internalized parasites (Fig. 1B). This was confirmed by orthogonal and single-stack confocal images, which revealed parasites fully internalized and positioned in the perinuclear region of host cells (Figs. 1C–D).

**Fig. 1.**
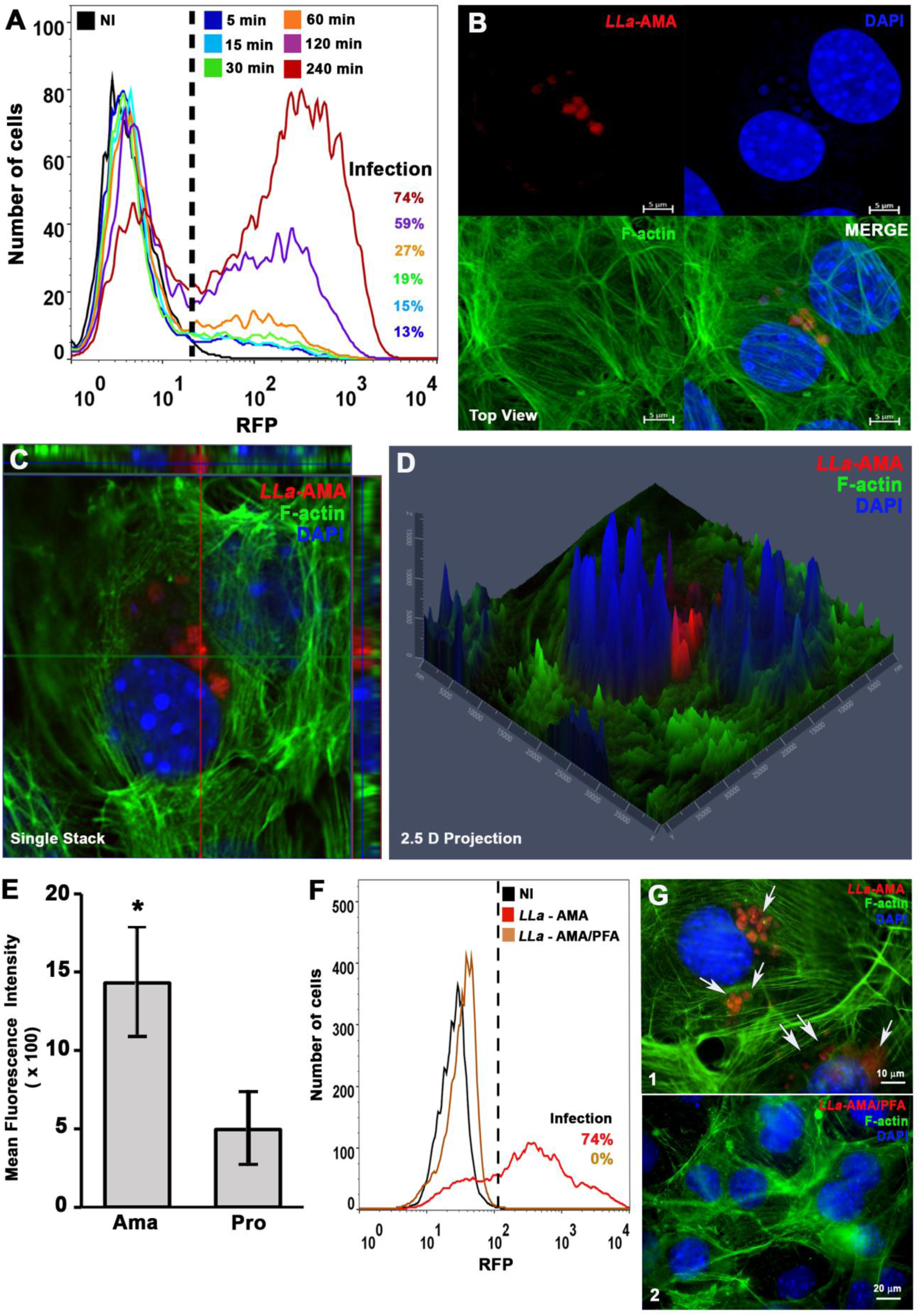
– *L.amazonensis* amastigotes induce invasion in MEFs and harbor the perinuclear region of the host cell. MEFs were incubated with *L. amazonensis* amastigotes and analyzed by FACS and fluorescence microscopy. **(A)** Time-course of MEF infection by *L. amazonensis* amastigotes analyzed by FACS. **(B)** Fluorescence microscopy analysis of MEF incubated for 4 h with *L. amazonensis* amastigotes (*LLa*-AMA, red) and stained by phalloidin-A488 (F-actin, green) to visualize host cell cytoskeleton and DAPI to stain nuclei (blue) – image obtained using Axio Imager ApoTome2 Microscope (Zeiss). **(C)** Orthogonal view revealing internalized amastigotes. **(D)** 2.5D of the infected cell. **(E)** *L. amazonensis* amastigotes are more infective to MEFs than promastigotes. After 4h infection with *L. amazonensis* amastigotes or promastigotes, cells were analyzed by FACS and the mean fluorescence intensity was used to compare the infectiveness of parasites to MEFs. The data represent the mean ± s.e.m. of three independent biological replicates * P = ≤ 0.05, Student′s t-test. **(F** and **G)** Infection of MEF by living (*LLa* – AMA) or PFA-fixed (*LLa* – AMA/PFA) *L. amazonensis* amastigotes as analyzed by **(F)** FACS and by **(G)** fluorescence microscopy (same labelling as in B). The images show that live parasites **(G-1)** are internalized into the cells (indicated by white arrows), residing near the nuclei (blue), whilst PFA-fixed parasites **(G –2)** were not observed inside the cells. Image obtained using a BX60 upright compound fluorescence microscope (Olympus).

Since we have previously demonstrated that the flagellated promastigote forms – *Leishmania*’s infective stage transmitted during the insect vector’s bite – can also invade MEFs (24), we compared the infection rates of promastigotes and amastigotes in these cells. Our results show that amastigotes were markedly more infective to MEFs than promastigotes (Fig. 1E), underscoring their enhanced ability to efficiently invade non-phagocytic cells. This property was also observed in other cell types, including muscle and epithelial cells (not shown), suggesting that the amastigote stage has evolved specific mechanisms for the invasion of non-phagocytic cells, supporting its role as infection-spreading agent within mammalian hosts. The invasion of non-phagocytic cells by amastigotes is dependent on parasite viability since PFA-fixed parasites were not detected inside MEFs, as assessed by flow cytometry (Fig. 1F) or F-actin labeling fluorescence microscopy (Fig. 1G) after contact of *LLa*-RFP.

Since *Leishmania* spp. typically reside within intracellular vacuoles that fuse with lysosomes along the endocytic pathway, we also investigated whether amastigotes were enclosed within phagolysosomes. Infected cells were labeled with anti-LAMP-1 and secondarily probed to visualize host cell lysosomes (green), along with phalloidin (magenta) to highlight the host cell F-actin cytoskeleton. Samples were then analyzed using ultra-resolution confocal fluorescence microscopy (Figs. 2A – 1 and 2). In Fig. 2A-2, with blue and green filters only, the individual round-shaped phagolysosomal compartments that surround amastigotes are clearly visible in the perinuclear region of the host cell.

**Fig. 2.**
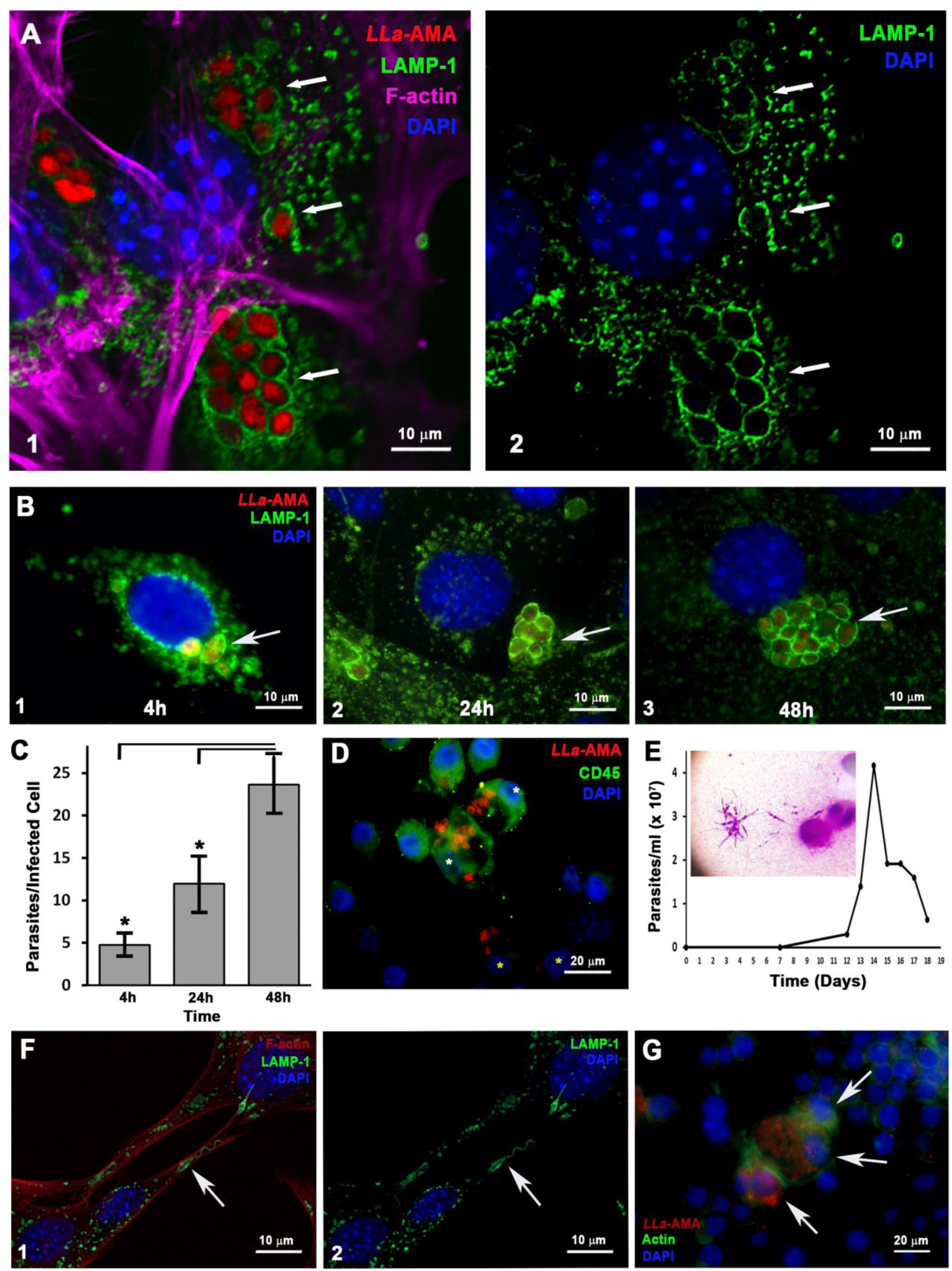
– *L. amazonensis* amastigotes reside within intracellular vacuoles enriched in lysosomal markers that support parasite replication and viability in MEFs. (A 1 and 2) Fluorescence microscopy analysis of MEF infected by *L. amazonensis* amastigotes (*LLa*-AMA, red) and stained by anti LAMP1 antibody to visualize host cell lysosomes (LAMP1, green), phalloidin-A633 (F-actin, magenta) to visualize host cell cytoskeleton and DAPI to stain nuclei (blue). **(A2)** Image showing only green (LAMP1-lysosomes) and blue (DAPI-nuclei) channels to highlight the parasitophorous vacuoles rich in lysosomal markers. Images were taken using Airyscan technology in a LSM 880 microscope (ZEISS) to obtain super resolution images. **(B)** To evaluate whether parasites were able to replicate within MEFs, the cells were infected with *L. amazonensis* amastigotes (*LLa*-AMA, red), fixed with PFA at the indicated time points, stained as in A and analyzed. The results show the presence of intracellular amastigotes close to host cell nuclei after 4h **(B-1)**, 24h **(B-2)** and 48 **(B-3)** of infection, living in individual vacuoles rich in lysosomal markers (white arrows). **(C)** The number of parasites per infected cell was obtained by manual counting after 4, 24 and 48h hours of infection. The data represent the mean ± s.e.m. (n = 25) * P = ≤ 0.05, Student′s t-test. **(D)** Fluorescence microscopy analysis of infected MEFs co-cultured with RAW cells (macrophage-like cells) for 48 hours. Cells were fixed with PFA and labeled with CD45 (green) to distinguish RAW cells, and DAPI to visualize nuclei (blue). Images were captured using the BX60 microscope (Olympus). RAW cells become infected (white asterisks) upon contact with infected MEFs (yellow asterisks). **(E)** Flagelated promastigote forms appeared in cell culture (inserted image) after infected MEFs were scrapped from the culture dish, inoculated in insect media M199 and cultivated at 24°C – the graph represents the growth curve of the re-transformed parasite. **(F)** Fluorescence microscopy analysis of MEFs infected with *L. amazonensis* promastigotes re-transformed from intracellular amastigotes. Cells were labelled with anti-LAMP1 antibody to visualize host cell lysosomes (LAMP1, green), phalloidin-A633 (F-actin, red) to visualize the host cell cytoskeleton, and DAPI to stain the nuclei (blue). The image demonstrates parasites surrounded by a tight lysosome coat (green, white arrows) acquired during their internalization into MEF cells. **(G)** RAW cells infected with *L. amazonensis* promastigotes re-transformed from intracellular amastigotes, cells were labeled with phalloidin-A488 (green) to visualize host cell cytoskeleton and DAPI (blue) to visualize nuclei. Images were captured using the Axio Imager ApoTome2 Microscope (Zeiss).

### *L. amazonensis* amastigotes replicate and persist within MEFs and are able to re-transform into promastigotes

To determine whether amastigotes internalized by non-phagocytic cells completed their intracellular stage following invasion, we monitored their development over time. Our results demonstrate that the parasites were able to replicate, both themselves and their individual vacuoles, within the first 48 h after invasion (Fig. 2B – 1 to 3). The number of parasites per infected cell roughly doubled each day, from 4 to 48 h (Fig. 2C), clearly indicating parasite proliferation. When MEFs infected for 48 h were co-cultured with RAW cells and labeled with anti-CD45 to selectively highlight the macrophage-like cells, amastigotes were observed spreading to the RAW cell population (Fig. 2D), establishing intracellular infection, showing that parasites were not only viable but also infective. In a parallel experiment, after 72 hours of infection – during which parasites persisted and multiplied as typical intracellular amastigotes – MEFs were detached by scraping, transferred to insect medium, and incubated at 24 °C to assess their reversion to promastigotes. Our results show that flagellated promastigotes appeared as host cells died, and the resurgent extracellular promastigotes successfully multiplied in insect media (Fig. 2E). These re-transformed promastigotes were also capable of infecting MEFs (Fig. 2F) and macrophages (Fig. 2G) in vitro, maintaining their ability to re-transform, proliferate, and infect different cell types, thereby preserving their plasticity in response to each host environment. Preliminary results suggest that promastigotes derived from intracellular amastigotes that were recovered from MEFs exhibit even increased infectivity compared to their original counterparts (data not shown). Taken together, our data show that the parasite not only survives and proliferates within MEFs but also preserves their infectivity, either by establishing new infections in host cells or by resuming development of promastigotes which are the stages capable of establishing sandfly infections after parasitized cells are taken up by the vectors during a blood meal.

### Invasion of MEFs by *L. amazonensis* amastigotes involves recruitment of host cell F-actin and subsequent fusion of the parasitophorous vacuole with host cell lysosomes

Since both F-actin and host cell lysosomes are known to play crucial roles in the invasion of host cells by various intracellular parasites – and given that host cell lysosomes were previously shown to be involved in the early stages of *L. amazonensis* promastigote invasion in MEFs (24) – we sought to investigate the involvement of the host actin cytoskeleton and lysosomes in the invasion of MEFs by amastigotes. To this end, we performed infection assays and analyzed cells at early time points after inoculation of parasites into host cell cultures to evaluate the invasion process. Cells were stained with phalloidin to visualize F-actin (green) and with anti-LAMP1 antibodies (magenta) to detect host lysosomes (Fig. 3). The results revealed that F-actin, but not lysosomes, is recruited to the sites of amastigote invasion (Fig. 3B–E). A prominent ring-like structure of F-actin was observed surrounding invading amastigotes (Fig. 3A–B, Fig. S4A–C), while, as expected, no F-actin recruitment was observed at promastigote entry sites (Fig. S4D). Further fluorescence microscopy analysis of an infection time-course confirmed that amastigotes trigger intense F-actin polymerization during the early stages of invasion (Fig. 4A), with parasites appearing surrounded by a dense F-actin network as they enter host cells (Fig. 4A–B, white arrows). At later time points, amastigotes were fully internalized within vacuoles that no longer exhibited F-actin staining (Fig. 4C, white arrows). On the other hand, recently internalized parasites did not co-localize with LAMP1 (Fig. 4D), a feature that became evident only at later time points following internalization (Fig. 4E–F). Quantitative analysis of F-actin and LAMP1 surrounding the parasites over time confirmed a clear temporal distinction: F-actin was associated with early parasitophorous vacuoles, whereas LAMP1 predominantly marked mature vacuoles (Fig. 4G). This pattern sharply contrasts with what we have previously observed during promastigote invasion, where rapid lysosome mobilization to the cell periphery results in early fusion with nascent parasitophorous vacuoles, forming a dense LAMP1-positive coat (24) (also evident here in Fig. 2F and Fig. 3F).

**Fig. 3.**
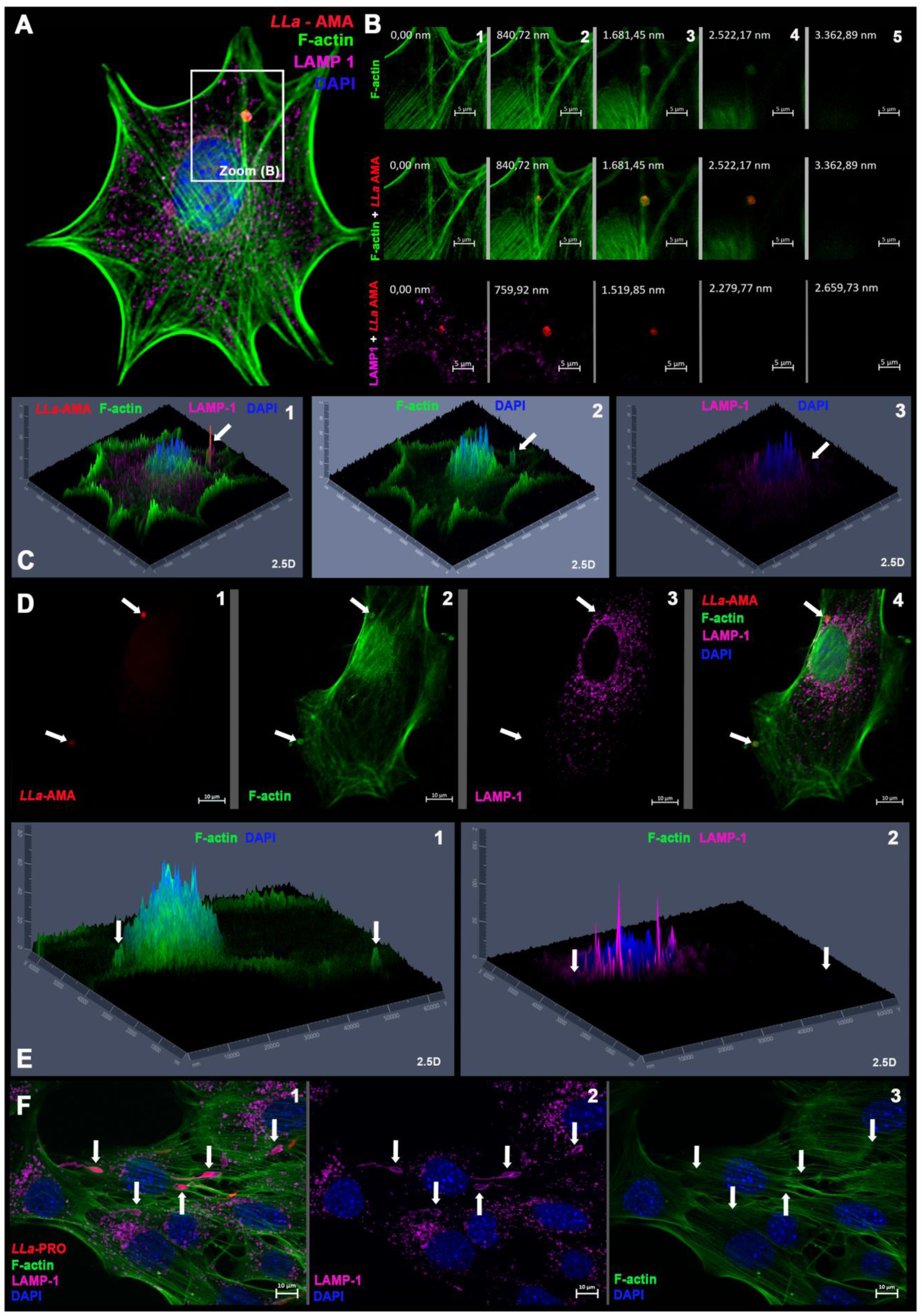
– *L. amazonensis* amastigotes trigger localized actin polymerization at invasion sites during cell invasion in MEFs. MEFs were incubated with *L. amazonensis* amastigotes (*LLa*-AMA, red) for 60 minutes. Cells were stained with phalloidin-A488 to visualize host cell F-actin (green), anti-LAMP1 antibody to visualize host cell lysosomes (magenta), and DAPI to visualize nuclei (blue). **(A)** An amastigote form (*LLa*-AMA, red) invading a MEF triggering local actin polymerization at the invasion site. **(B)** z stacks of the zoomed-in area indicated in panel A, where 1 to 5 represent single stacks, from the bottom to the top, in different channels, showing that the amastigote invades the host cell by an actin-dependent and lysosome-independent manner. **(C)** 2.5D projection of the stack represented in B3 showing different channels (1 to 3, as indicated). Amastigote (*LLa*-AMA, red) invading (white arrow) a MEF by recruiting (2) F-actin (green) and not (3) lysosomes (magenta). **(D)** This phenomenon was consistently observed in cells being infected by amastigotes. This image shows two distinct amastigotes entering the same cell (invasion sites pointed by white arrows). (**1**) Amastigotes (*LLa*-AMA-white arrows), (**2**) F-actin nests (green) are formed at invasion sites that exclude (**3**) lysosomes, and (**4**) merged image. **(E)** 2.5D projection of panel D following the indicated channels. **(F)** Same experiment, but using *L. amazonensis* promastigotes (*LLa*-PRO, red), showing that actin polymerization (F-actin) is not induced during promastigote entry in MEF and that, instead, parasites are enwrapped by a tight coat of lysosomes (magenta) that perfectly delineate the parasite (white arrows). Images were captured using the Axio Imager ApoTome2 Microscope (Zeiss).

**Fig. 4.**
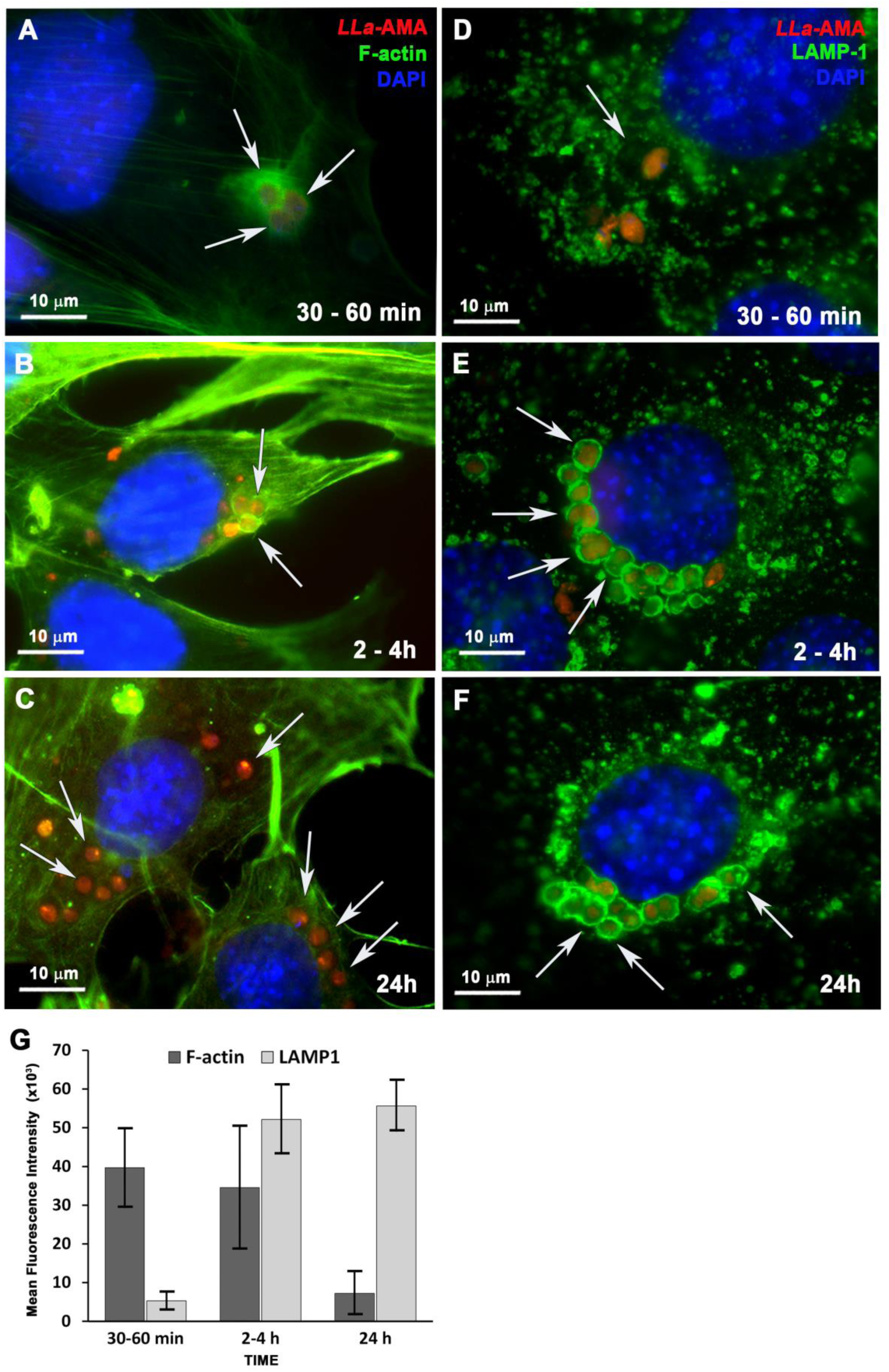
– Cell invasion by *L. amazonensis* amastigotes in MEF involves early recruitment of host cell actin and subsequent fusion of the parasite-containing endosome with host cell lysosomes. MEFs were infected with *L. amazonensis* amastigotes (*LLa*-AMA, red), fixed with PFA at the indicated time points, stained with phalloidin-A488 to visualize host cell F-actin (green) or anti-LAMP1 antibody (green) to visualize host cell lysosomes and with DAPI to visualize the nuclei (blue). **(A and B)** Recently internalized amastigotes (*LLa*-AMA) are surrounded by actin filaments (F-actin). **(C)** At late time points after cell invasion no actin nest is observed surrounding the internalized parasites. **(D)** recently internalized parasites do not show lysosomal markers surrounding the parasitophorous vacuoles (LAMP1, green). **(E and F)** Following cell invasion, the amastigotes inhabit intracellular vacuoles that intensely fuse with host cell lysosomes (LAMP1, green) and surround host cell nuclei. **(G)** Quantification of F-actin and LAMP1 associated to *L. amazonensis* containing-vacuoles during amastigote infection in MEFs. The fluorescence intensity of phalloidin-A488 (F-actin) or LAMP1 (lysosomes) associated to parasite vacuoles were measured for each time point using 25 individual vacuoles, the data represent the mean ± s.e.m. All images were acquired using a BX60 upright compound fluorescence microscope (Olympus). White arrows point to parasites and their vacuoles.

### *L. amazonensis* amastigotes induce intense membrane ruffling in MEFs during cell entry

To visualize the dynamics of the host cell plasma membrane during amastigote invasion, we performed SEM analyses of MEFs during cell invasion. The images revealed intense membrane ruffling in MEFs upon amastigote contact (Fig. 5A–C), but not during interaction with promastigotes (Fig. 5D–F). As previously demonstrated (24), promastigotes were observed entering MEFs through their flagella (Fig. 5F), during which the plasma membrane behaves differently, remaining flat and showing no ruffling. The invasion of MEFs by amastigotes clearly involved significant activity in the host cell plasma membrane, which wrapped around and ultimately enveloped the parasites (Fig. 5 B, C and H). These dramatic changes and the intense ruffling in the host cell plasma membrane were only observed at early moments of interaction with amastigotes, no longer present when after internalization of (Fig. 5G-I).

**Fig. 5.**
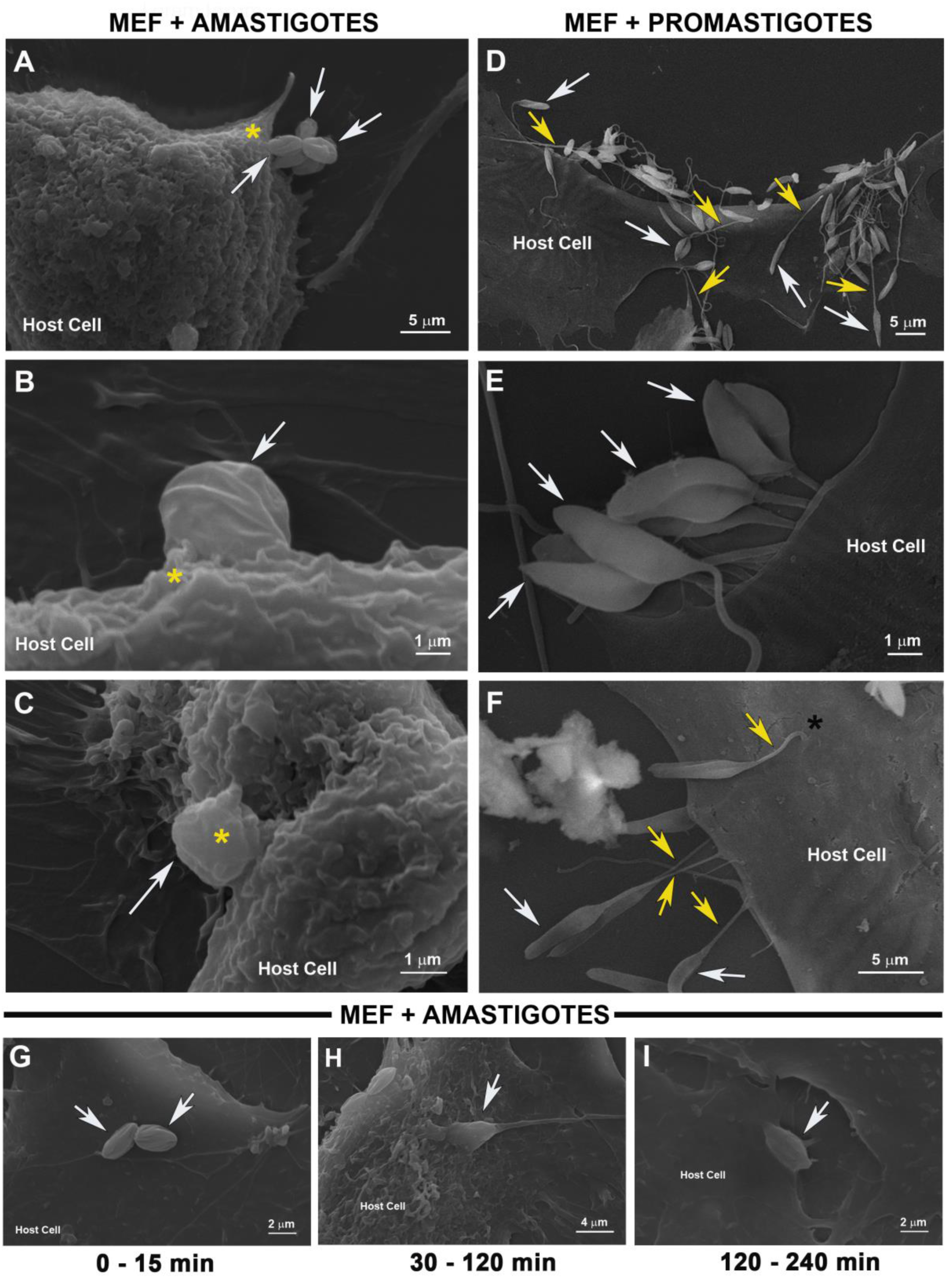
– *L. amazonensis* amastigotes but not promastigotes induce intense membrane ruffling in MEFs during cell invasion. *L. amazonensis* amastigotes and promastigotes were incubated with MEF for 30 min, after infection cells were fixed and prepared for scanning electron microscopy (MEV). **(A)** amastigotes (white arrows) interact with the host cell inducing an intense membrane ruffling (yellow asterisks). **(B and C)** Once bound to MEFs the parasites are gradually enwrapped by host cell plasma membrane leading to its internalization. **(D to F)** promastigotes (white arrows) strongly bind to host cells by their flagella (yellow arrows) forcing its internalization without provoking any membrane ruffling and invades MEFs by inserting themselves into the host cell (black asterisk in F). **(G to I)** Time course of cell invasion by *L. amazonensis* amastigotes (white arrows) in MEFs, as analyzed by MEV.

### *L. amazonensis* amastigotes depend on host cell actin polymerization to invade MEFs

Altogether, our data indicated that amastigotes are highly infective to fibroblasts with their entry involving host cell F-actin, in stark contrast to the actin-independent behavior observed during promastigotes infection of the same cell type. To confirm the requirement of host F-actin for amastigote entry into MEFs, we pre-treated cells with cytochalasin D to inhibit actin polymerization, followed by incubation with amastigotes or promastigotes. Our results showed that inhibition of F-actin polymerization completely blocked amastigote entry into MEFs (Fig. 6A, C). In contrast, promastigote infection of cytochalasin-D-treated MEFs is not blocked. On the contrary, infection increased approximately threefold compared to untreated controls (Fig. 6B, C). These contrasting findings are readily observed in the fluorescence microscopy images (Fig. 6D-I and Fig. S5), in which we can see that amastigotes stayed in the extracellular milieu of cytochalasin D-treated MEFs, unable to invade host cells (Fig. 6F,H), as opposed to untreated cells, where they entered encircled by F-actin (Fig. 6D) while promastigotes successfully entered cytochalasin D-treated MEFs (Fig. 6G) in vacuoles containing the lysosomal marker LAMP1 (Fig. 6I). Promastigote infection under actin-disrupting conditions resulted in both higher infection rates and an increased parasite burden per cell, in confirmation of our previous study (24). These findings were further validated using non-axenic amastigotes directly obtained from infected macrophages (Fig. S6) and extended to other non-phagocytic host cell types including Huvec, HepG2, and HeLa cells (Fig. S7), demonstrating that this phenomenon is not restricted to parasite from axenic cultures or to fibroblasts.

**Fig. 6.**
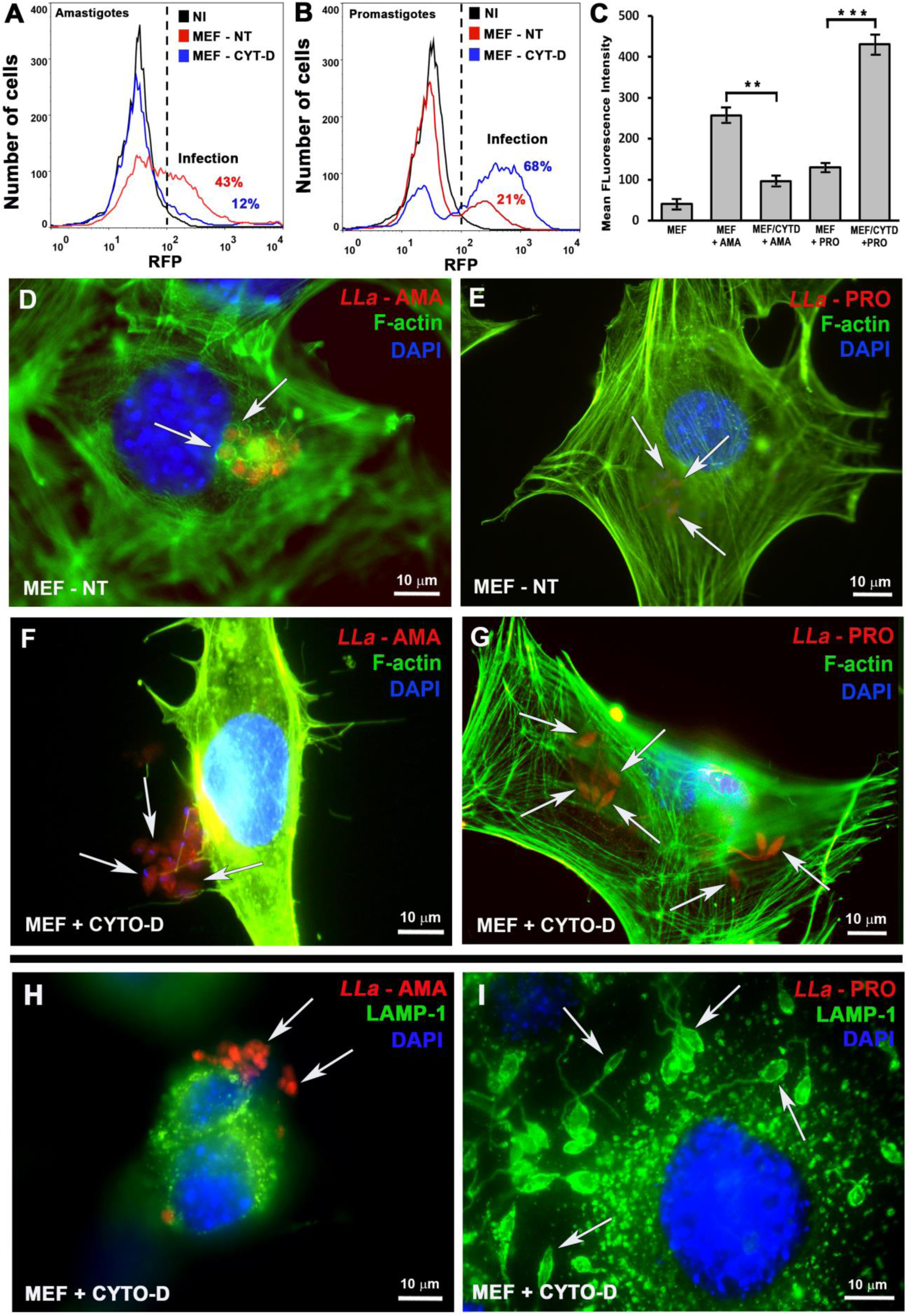
– Host cell F-actin drives *L. amazonensis* amastigote invasion in MEFs. **(A)** amastigotes and **(B)** promastigotes of *L. amazonensis* were incubated with MEFs with (red) or without (blue) cytochalasin-D pre-treatment to block actin polymerization and the cell population was analyzed by means of FACS to quantify infection. **(C)** Infection quantification in MEFs pre-treated or not with cytochalasin D and infected with either amastigotes or promastigotes. Infection rates were assessed based on the mean fluorescence intensity (MFI) of each cell population/treatment, as determined by flow cytometry (FACS). The data represent the mean ± s.e.m. of three independent biological replicates **P=0.007, ***P=0,001, Student′s *t*-test. **(D-G)** Cells from the experiments shown in A and B, infected with *L. amazonensis* amastigotes (*LLa* – AMA, red) or promastigotes (*LLa* – PRO, red); **(D and E)** untreated and **(F-G)** cytochalasin-D-treated cells were fixed with PFA, stained with phalloidin-A488 to visualize host cell F-actin (green) and DAPI to visualize the nuclei (blue). **(H and I)** same as F and G but cells were stained with anti LAMP1 (green) to visualize host cells lysosomes – all images were obtained using a BX60 upright compound fluorescence microscope (Olympus).

### RhoA and Rac1, are recruited to amastigote invasion sites and invasion is drastically impaired by ROCK inhibition

Based on our previous findings that host cell actin plays a pivotal role in amastigote invasion of non-phagocytic cells, we next investigated whether members of the Rho GTPase family, key regulators of actin polymerization, are recruited to parasite entry sites. To this end, we performed infection assays and stained cells with antibodies against RhoA and Rac1, followed by fluorescence microscopy analysis. Both RhoA and Rac1 were found to co-localize with invading amastigotes (Fig. 7), where a strong enrichment at actin-rich invasion sites was evident (Fig. 7, red arrows). This co-localization was markedly reduced in parasites located deeper within the cytoplasm, near host cell nuclei (Fig. 7A-B, white arrows), consistent with a more advanced stage of invasion. Since Cdc42, another member of the Rho GTPase family, has been reported to participate in the invasion of RAW cells by *T. cruzi* amastigotes (32), we also examined its recruitment in our model; however, only negligible Cdc42 staining was occasionally observed (data not shown). Additionally, in a complementary experiment, simultaneous labeling of RhoA and Rac1 demonstrated that both GTPases co-localize at the same parasite entry sites (Fig. 7C-G, white arrows, Fig. S9), suggesting that their coordinated activity contributes to the internalization process. Accordingly, a ROCK inhibitor – which selectively blocks ROCK activity downstream of RhoA – caused a reduction of nearly 70% and 50% in infection rates and in the number of intracellular parasites per cell, respectively (Fig. 8). Together, these findings demonstrate the critical role of RhoA and probably Rac1 in promoting amastigote entry into fibroblasts, by fueling host cell actin polymerization.

**Fig. 7.**
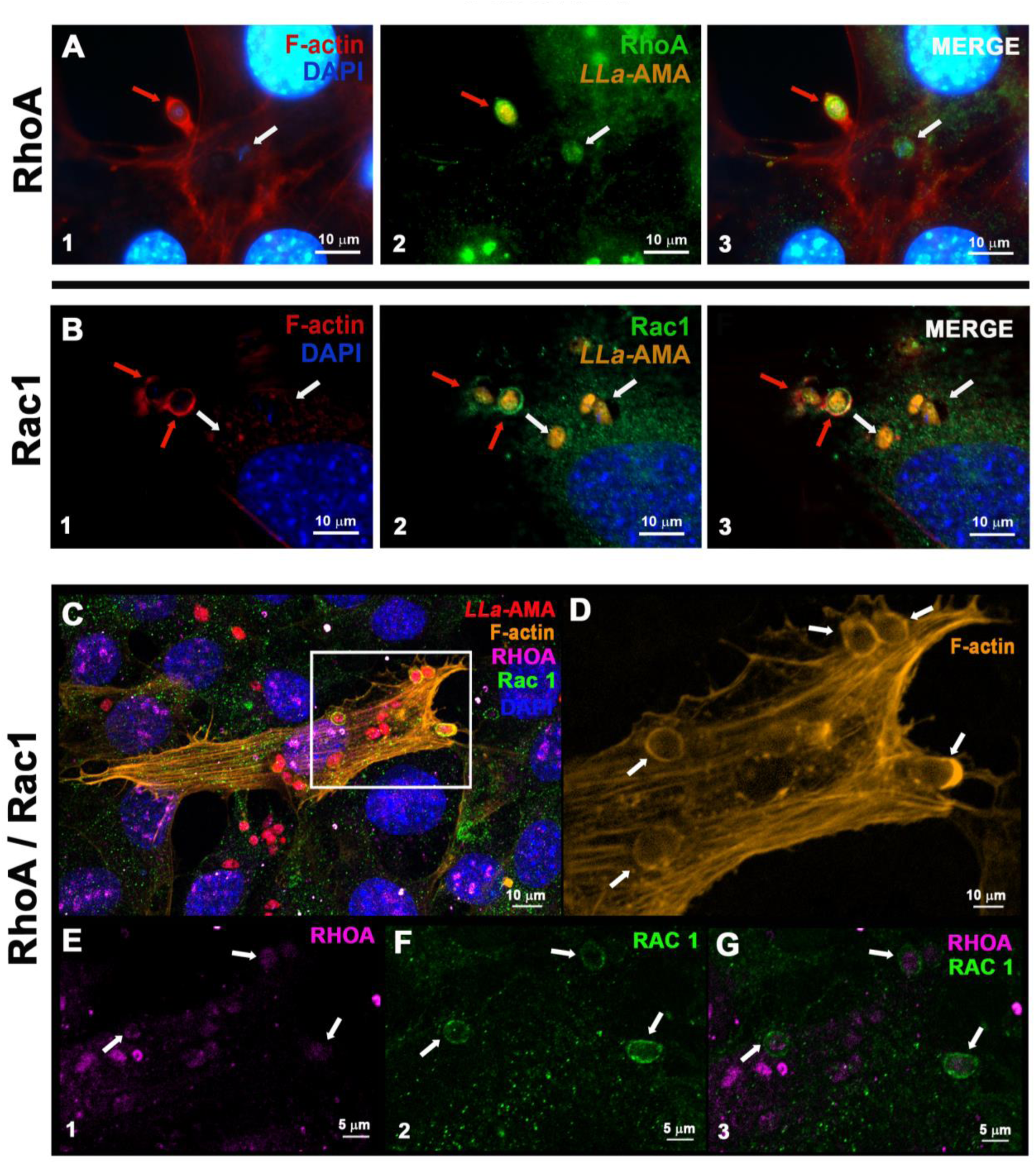
– The proteins of Rho GTPase family RhoA and RAC1 are recruited to *L. amazonensis* amastigotes invasion foci in MEFs. (**A-B**) MEFs were incubated with *L. amazonensis* amastigotes (*LLa*-AMA, orange) for 60 minutes. Cells were stained with phalloidin-Alexa Fluor 633 to visualize host cell F-actin (red), anti-RhoA or anti-Rac1 antibodies to detect the presence of these GTPases (green), and DAPI (blue) to label nuclei. (**A1**-**B1)** show actin polymerization at sites of parasite entry (red arrows), which co-localizes with **(A2)** RhoA and **(B2)** Rac1. White arrows indicate parasites at later stages of infection, residing in vacuoles that lack F-actin accumulation and exhibit weaker labeling for both GTPases. **(A3 and B3)** show the merged images. Images were taken using confocal LSM 880 microscope (ZEISS). **(C-G)** RAC1 and RhoA are both recruited to F-actin-rich sites during the invasion of MEFs by amastigotes. **(C)** Fluorescence microscopy analysis of MEFs infected for 60 minutes with *L. amazonensis* amastigotes (*LLa*-AMA, red), labelled with anti-Rac1 (green) and anti-RhoA (magenta) antibodies to assess the recruitment of both proteins to sites of parasite entry. F-actin was labeled with phalloidin-A555 (orange), and nuclei were stained with DAPI (blue). **(D)** Magnified view of the region highlighted in panel C showing F-actin accumulations displaying a round-shaped organization at sites of amastigote invasion (white arrows). **(E-G)** Individual fluorescence channels showing labeling of **(E)** RhoA, **(F)** RAC1, and **(G)** the merged image showing the co-localization. Images were acquired using the spectral detector ChS1 on an confocal LSM 880 microscope (ZEISS).

**Fig. 8.**
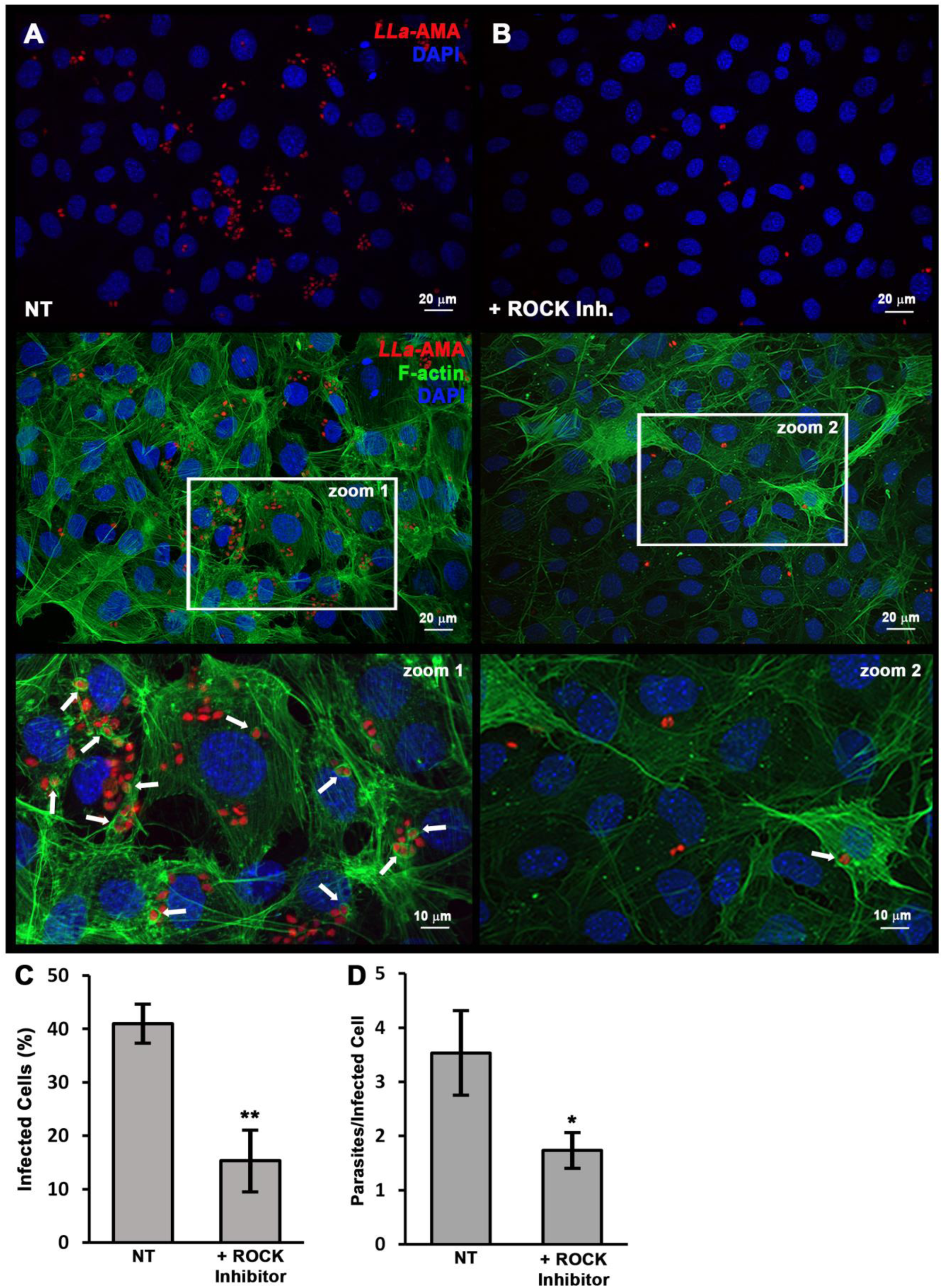
– Inhibition of Rho GTPases reduces the entry of *L. amazonensis* amastigotes in MEFs. MEF monolayers were pre-treated with a ROCK inhibitor for 60 minutes, washed, and then incubated with *L. amazonensis* amastigotes. After 60 minutes of infection, cells were fixed and stained with phalloidin-A488 (green) and DAPI (blue) to label F-actin and nuclei, respectively. **(A)** Untreated MEFs exhibited a higher number of internalized amastigotes (*LLa*-AMA, red) compared to **(B)** MEFs pre-treated with ROCK inhibitor. Lower panels show phalloidin-A488 staining (F-actin, green), revealing that most amastigotes were internalized in untreated cells, whereas ROCK inhibitor-treated cells displayed reduced internalization and diminished F-actin accumulation at invasion sites. Enlarged images (zoom 1 and zoom 2) highlight the presence of F-actin-rich structures (white arrows) surrounding invading amastigotes in untreated cells, which were less prominent following ROCK inhibition. These qualitative observations are supported by quantitative analyses showing **(C)** the infection rate and **(D)** the average number of parasites per infected cell. Data represents the mean ± s.e.m. from three independent biological replicates *P = ≤ 0.05, **P= ≤ 0.01, Student′s t-test. Images were captured using the Axio Imager ApoTome2 Microscope (Zeiss).

## DISCUSSION

Although numerous studies over the years have reported intracellular stages of *Leishmania* spp. within non-phagocytic cells in vivo and in vitro (8–18,20,22,24,33–36), the persistent and often misleading notion that both promastigotes and amastigotes of *Leishmania* spp. must be phagocytosed to infect cells still prevails. Moreover, the contribution of parasitized non-phagocytic cells to infection progression, along with the mechanisms underlying their invasion by *Leishmania* spp., remains largely speculative and underexplored. Until recently, little was known about how non-phagocytic cells could be invaded by the two infective forms of this parasite. In previous work we have demonstrated that promastigotes – the infective form transmitted through the insect vector’s bite – invade non-phagocytic cells by subverting a common membrane repair mechanism triggered by a parasite-induced membrane lesion. The entry of Ca^2+^ triggers the recruitment and focal exocytosis of lysosomes at the site of cell invasion, which supplies the endomembranes that cause invagination of a nascent parasitophorous vacuole, culminating in parasite internalization (24). This mechanism appears to be evolutionarily conserved among intracellular trypanosomatids since it is also observed in *Trypanosoma cruzi* (25). Here, we demonstrate that *L. amazonensis* amastigotes – the infective form responsible for parasite replication, tissue dissemination and disease development, can also efficiently invade non-phagocytic cells (Fig. 1 and Fig. S7), albeit through mechanism fundamentally different from that employed by promastigotes. They are ultimately localized within parasitophorous vacuoles, which subsequently fuse with lysosomes along the endosomal pathway where they survive and replicate (Fig. 2A-B and Fig. 4E-F). Cell invasion is directly induced only by live parasites, as PFA-fixed parasites were not detected inside host cells (Fig. 1F-G). This suggests that invasion is induced by the parasite, depending either on the integrity of surface protein ligands, or the secretion of parasite-derived virulence factors that facilitate entry, or both. Importantly, when infected cells were removed from the culture dish and transferred to insect medium, amastigotes re-differentiated into promastigotes and remained infective (Fig. 2E-F). This shows that parasites replicating inside non-phagocytic cells remain viable and retain all the machinery to differentiate into promastigotes. These findings suggest that infected non-phagocytic cells may serve as reservoirs for the vector. Notably, dogs, a primary reservoir of *L. infantum*, develop high parasite burdens throughout their skin, including within fibroblasts (16). Because *Leishmania* amastigotes accumulate within tissue-resident cells rather than circulating in the bloodstream, transmission to the vector occurs primarily when it ingests these infected host cells during blood meals. Thus, in addition to macrophages, other cell types can harbor amastigotes and directly contribute to vector infection, likely entering these cells via the mechanisms demonstrated in this study.

Beyond enabling unambiguous visualization of internalized parasites, labeling of the host cell F-actin cytoskeleton revealed that the early stages of invasion are characterized by intense actin polymerization localized at the host cell/parasite membranes interface (Fig. 3A-E, Fig. 4A-B, Fig.6D, Fig. 7A, B and D, Fig. S4 and Fig. S8 and Fig. S9), resembling the invasion mechanisms of intracellular bacteria such as *Rickettsia*, *Chlamydia*, *Shigella*, and *Listeria* (37,38). This F-actin–rich network enveloped newly internalized amastigotes (Fig. 3A-E) – an arrangement not observed during promastigote invasion of the same cell type (Fig. 3F and Fig. S4D). Notably, F-actin recruitment is a hallmark of non-phagocytic cell invasion by many intracellular pathogens (37–42) and, while central to phagocytosis, can also be subverted by pathogens to facilitate entry into non-phagocytic host cells (37–43). In the case of *Trypanosoma cruzi*, the intracellular protozoan that causes Chagas disease, which is phylogenetically related to *Leishmania* spp., it was observed that amastigotes attach to microvilli on the surface of HeLa cells, leading to the formation of a phagocytic cup-like structure under the parasite, which culminates in cell entry within 15-30 minutes of interaction (44,45). A recent report also suggests an important role for the host cell F-actin cytoskeleton in the process of cell invasion by both *T. cruzi* trypomastigotes and amastigotes, although pointing to a greater importance of F-actin recruitment when the infective form in question is the amastigote (46).

Upon contact with *L. amazonensis* amastigotes, host cells exhibit intense membrane ruffling (Fig. 5A-C), a response not observed following exposure to promastigotes (Fig. 5D-F), providing visual evidence that the membrane dynamics involved in the two invasion processes are indeed distinct. The strong inhibition of amastigote internalization upon treatment of cells with cytochalasin-D (Fig. 6, Fig. S6 and Fig. S7) demonstrated that F-actin polymerization is not simply associated with parasite entry but is essential for the process. Again, the distinction between amastigote and promastigote invasion is highlighted by the finding that disrupting host cell actin polymerization during promastigote entry actually increases the parasite-to-cell ratio compared with cells with intact F-actin (Fig. 6C, G, I). This is possibly due to the indirect effects of cytochalasin D, either causing lysosome redistribution to the cell periphery, or facilitating membrane invaginations, thereby facilitating promastigote entry, as we have previously demonstrated (24). Actin-independent membrane invagination with lysosome-derived endomembrane supply *versus* F-actin-driven membrane ruffling with outward projections of the plasma membrane seem to represent hallmark differences between amastigote and promastigote invasion processes, respectively (Fig. 5 C and F). Regardless of the infective form, the intracellular parasites are ultimately localized within parasitophorous vacuoles that fuse with lysosomes along the endosomal pathway, which typically cluster near the host cell nucleus (Fig. 2 A-B Fig. 4E-F and Fig. S6 B).

One significant finding of the present study was that among the three prototypical Rho GTPases, which are key regulators of actin rearrangement and polymerization within the host cell, RhoA and Rac1, but not Cdc42, are intensely recruited to the infection sites, co-localizing with the invading parasites (Fig. 7, Fig. S8 and Fig. S9). This suggests that the formation of filipodia (47) is not essential for the interiorization of amastigotes. In contrast, the recruitment of RhoA and Rac1 to the site of parasite-host cell contact suggests that these Rho GTPases and that the formation of lamellipodia and stress fibers (47,48) are required for the amastigote invasion process. Indeed, this seems to be the case at least for RhoA, since the inhibition of ROCK, a RhoA-activated kinase, drastically reduces the infection rate of MEFs by amastigotes (Fig. 8). As ROCK phosphorylates 1) LIMK-1 and LIMK-2, which, through inactivation of cofilin, help maintain F-actin network, 2) myosin light chain (MLC) leading to an increase in actomyosin contractility, 3) ERM proteins (ezrin, radixin, and moesin) and NHE1 (Na+/H+ –Exchanger 1) to enhance coupling of the actin cytoskeleton to integral membrane proteins (47) it is likely that one or more components of these ROCK-dependent pathways are required for amastigotes to invade non-phagocytic cells. In this regard, it is possible that the parasite has evolved the ability to trigger actin-dependent cell entry through multiple routes, ensuring a fast and efficient cell entry.

The internalization of amastigotes into non-phagocytic cells has previously been linked to Cdc42 recruitment in CHO cells, with F-actin surrounding the parasites (49). In contrast, in the MEF model studied here, Rac1 and RhoA, but not Cdc42, were clearly detected at sites of parasite entry (Fig. 7, Fig. S8 and Fig. S9). In the case of Rac1, it has been previously demonstrated that during invasion of professional phagocytes by *L. donovani* amastigotes, this is the principal GTPase involved (50). Taken together, our findings, along with previous reports (49,50), suggest that *Leishmania* amastigotes exploit multiple canonical actin-dependent pathways to ensure efficient and rapid invasion of host cells, including non-phagocytic ones, transiently endowing them with the ability to engulf such a large parasite. The specific effectors produced by the parasite to mediate this process remain unknown to date and likely represent important virulence factors.

Macrophages have long been known to become infected by *Leishmania* amastigotes through indirect routes, such as the phagocytosis of apoptotic bodies from previously infected cells. This mode of infection, known as the Trojan horse hypothesis, is thought to increase macrophage susceptibility to parasitism, since clearance of apoptotic bodies does not activate their microbicidal effector functions. In this context, non-phagocytic infected cells could also serve as carriers of amastigotes to macrophages, as these cells eventually undergo senescence or apoptotic cell death, further contributing to parasite dissemination. Supporting this notion, our co-culture experiments showed that phagocytic cells became infected when incubated with infected fibroblasts (Fig. 2D).

Importantly, although macrophages remain the predominant cell type infected with *Leishmania* in lesions – likely reflecting the prevalence of the Trojan horse mechanism – other pathways can also amplify infection by releasing amastigotes into the extracellular milieu (51). Exocytosis-like expulsion of parasites or their liberation following necrotic death of host cells can release free amastigotes (52–54). Based on our findings, these extracellular amastigotes may be capable of invading virtually any available host cell type in the absence of opsonization, using the mechanism we describe here. Importantly, this property suggests that amastigotes may also directly invade macrophages constantly attracted to lesion sites through routes that bypass conventional phagocytosis. Such alternative entry could allow parasites to establish infection in a ‘silent’ manner, minimizing macrophage activation and thereby evading antimicrobial responses. Of note, many non-phagocytic cell types that harbor parasites in vivo are long-lived and, unlike professional phagocytes, lack the capacity to mount effective immune responses, thereby potentially serving as safe reservoirs for parasite persistence. Furthermore, non-phagocytic cells may serve as protective niches that shield *Leishmania* from commonly used drugs such as meglumine antimoniate and amphotericin B, which primarily target infected macrophages (55). Parasites residing within these cells may thus evade both immune clearance and standard chemotherapeutic approaches, contributing to frequent relapses following spontaneous or treatment-induced lesion healing.

The ability of amastigotes to invade non-phagocytic cells may have broad implications for *Leishmania* biology and for disease pathogenesis. In addition to serving as reservoirs, the ability of *Leishmania* to invade non-phagocytic cells may be crucial for parasite dissemination and visceralization. Following inoculation by the sand fly vector, parasites must travel from the bite site to establish infection in various tissues as amastigotes. The ability to cross cellular barriers appears fundamental for *Leishmania*, as these parasites can spread from the site of the bite throughout the skin, leading to disseminated lesions (a feature observed in *L. amazonensis* infection) or reach internal organs, causing visceralization (a hallmark of *L. infantum* and *L. donovani* infections). Although the exact mechanisms of dissemination remain largely unknown, it is plausible to hypothesize that the ability of amastigotes to induce their entry into any cell via the mechanisms described here might enable them to traverse tissues and navigate host cellular layers and cross tissue barriers, a critical step that could facilitate systemic spread. Moreover, species-specific differences in this pattern of invasiveness may help explain why some *Leishmania* species cause localized skin lesions while others lead to disseminated cutaneous or visceral disease.

By looking beyond classical phagocytosis and understanding previously unrecognized invasion mechanisms, as well as identifying the virulence factors that enable *Leishmania* to invade virtually any cell type, we may not only gain a better understanding of alternative routes of infection but also uncover new strategies to block or control amastigote invasion and combat disease. Such discoveries could reveal novel drug targets, guide vaccine development, and ultimately transform approaches to controlling and treat leishmaniases. Fully elucidating the spectrum of cellular invasiveness displayed by the pathogen, including the role of non-professional phagocytes both as long-lived reservoirs that support parasite persistence and as barriers that amastigotes must cross to disseminate, is critical to understanding parasite persistence, spread, and disease progression.

## ACKNOWLEDGMENTS AND FINANCIAL SUPPORT

We would like to thank Dr. Norma Andrews, Dr. Santuza Teixeira, Dr. Leda Quercia Vieira and Dr. Juliana Menezes for their support and generosity during the realization of this project and Dr. David Sacks for kindly providing the RFP-expressing parasites used in this work. Dr. Norma Andrews kindly reviewed this manuscript. We also would like to thank CAPI-ICB-UFMG (Centro de Aquisição e Processamento de Imagens) for all support with imaging and the different microscopy techniques used in this work, and the Flow Cytometry Laboratory-ICB-UFMG for support with all FACS analysis. This work received support from CNPq (Conselho Nacional de Desenvolvimento Científico e Tecnológico), TC-G and MFH are CNPq fellow and TQ-O is a CAPES (Coordenação de Aperfeiçoamento de Pessoal de Nível Superior) fellow.

## COMPETING INTERESTS AND FINANCIAL DISCLOSURE

The authors declare no competing interest and the funders had no role in study design, data collection and analysis, decision to publish, or preparation of the manuscript.

## LEGEND TO FIGURES

**Fig. S1.**
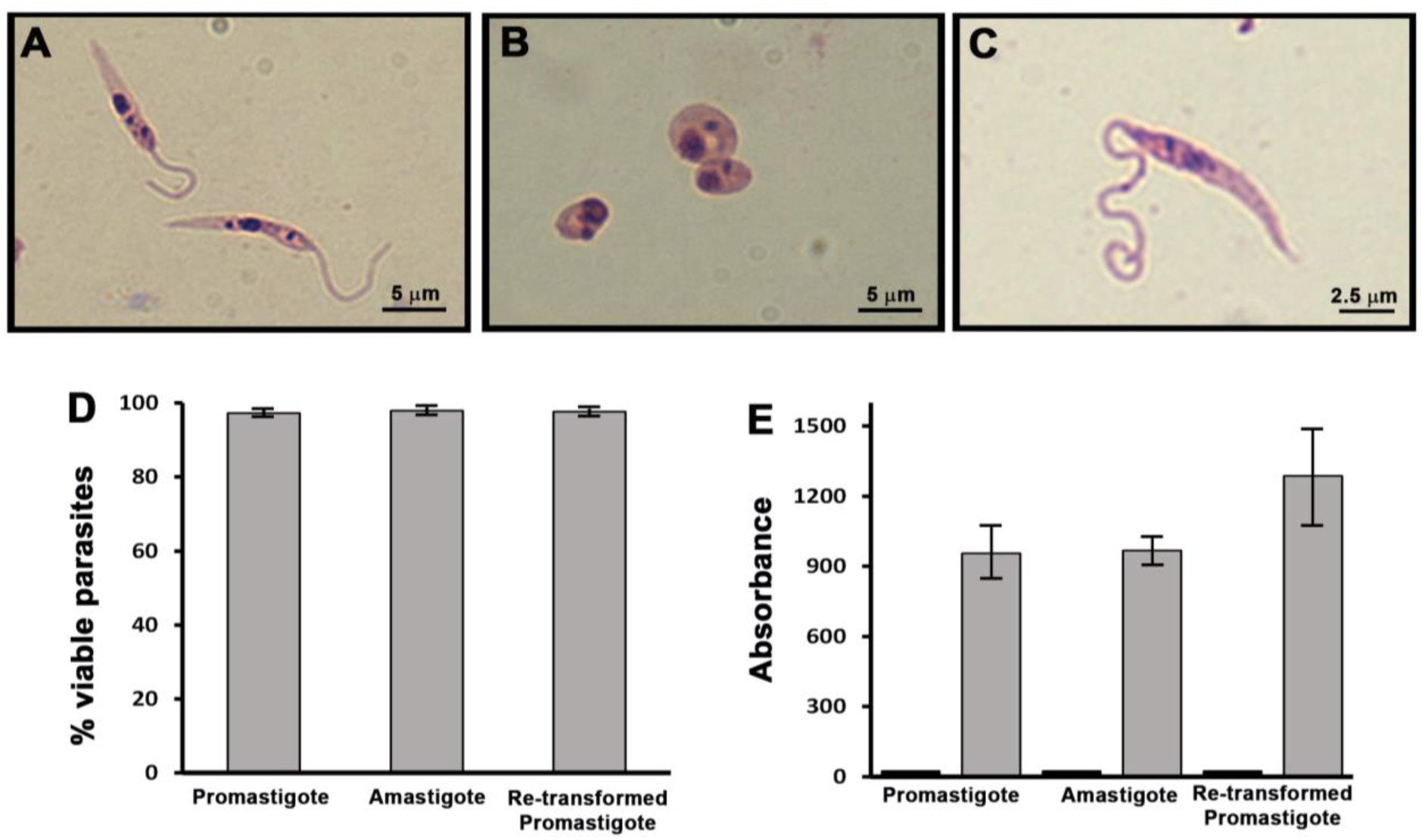
– *L. amazonensis* cultivation and obtention of axenic amastigotes. A, B, and C – Morphology of the parasites in their different evolutionary forms, before and after in vitro transformation. **(A)** axenic promastigotes, with apparent flagellum, nucleus and kinetoplast. **(B)** axenic rounded shaped amastigotes, without apparent flagellum, showing the nucleus and the kinetoplast. **(C)** axenic promastigotes re-transformed from the amastigote culture shown in B. **(D)** viability of promastigotes, amastigotes and re-transformed promastigotes, based on parasite counting in the presence of the vital dye erythrosine. **(E)** viability of promastigotes, amastigotes and re-transformed promastigotes using the MTT method. Dead parasites, fixed with PFA, were used as a negative control (black bars).

**Fig. S2.**
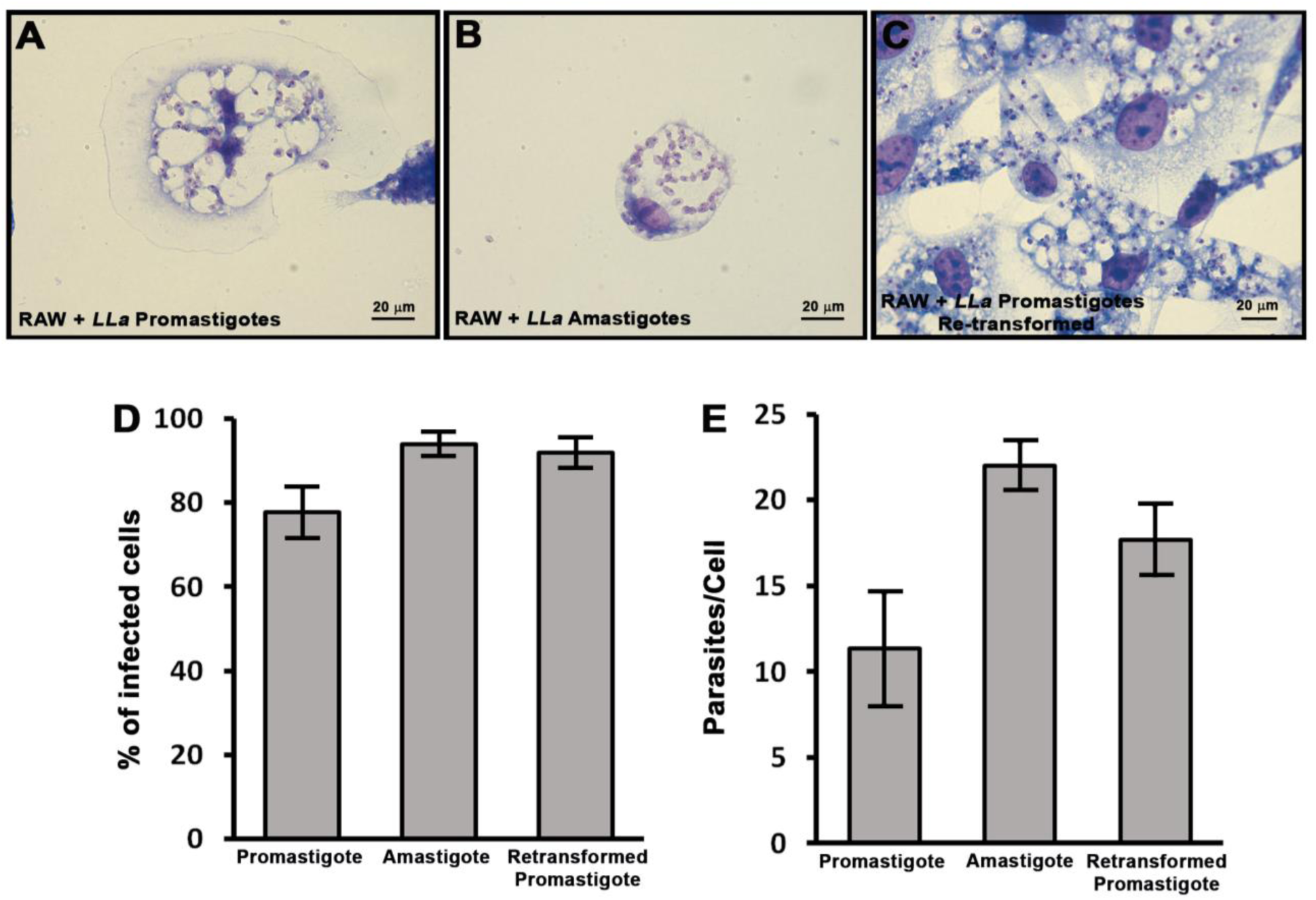
– Infection of macrophages with promastigotes, amastigotes, and re-transformed promastigote forms of *L. amazonensis*. Macrophages were infected with *L. amazonensis* (A) axenic promastigotes, **(B)** axenic amastigotes and **(C)** promastigotes re-transformed from axenic amastigotes for 4 hours at 37°C. After 24 hours, cells were fixed and stained using the panoptic method. **(D)** and **(E)** Quantification of infection and parasite load, confirming parasite infectivity for all evolutionary forms used. All images were acquired using a BX60 upright compound fluorescence microscope (Olympus).

**Fig. S3.**
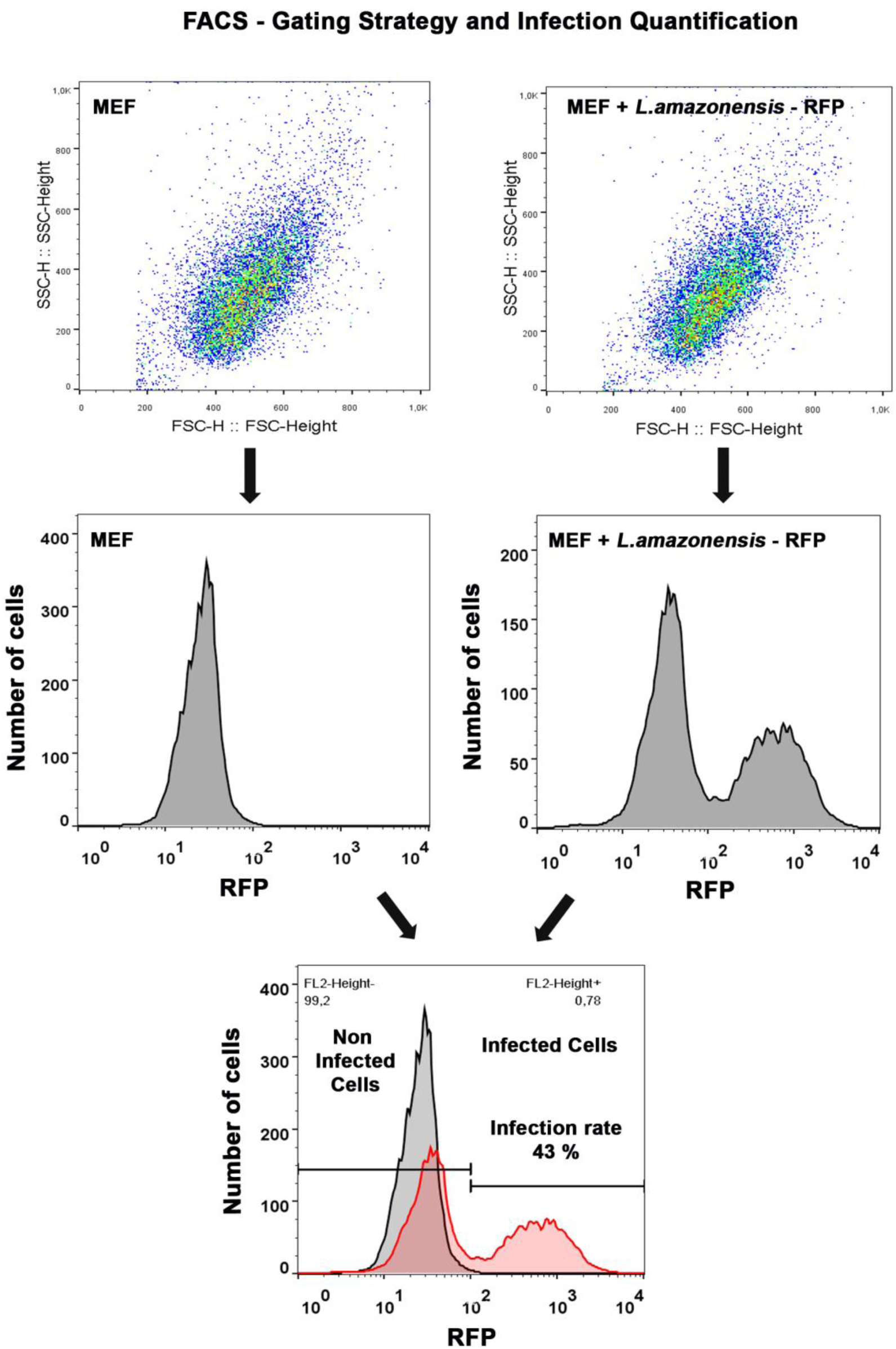
– Gating strategy used for the quantification of MEF infection by *L. amazonensis-*RFP by FACS. Schematic figure showing how flow cytometry data were acquired and processed from acquisition to the obtention of infection rates.

**Fig. S4.**
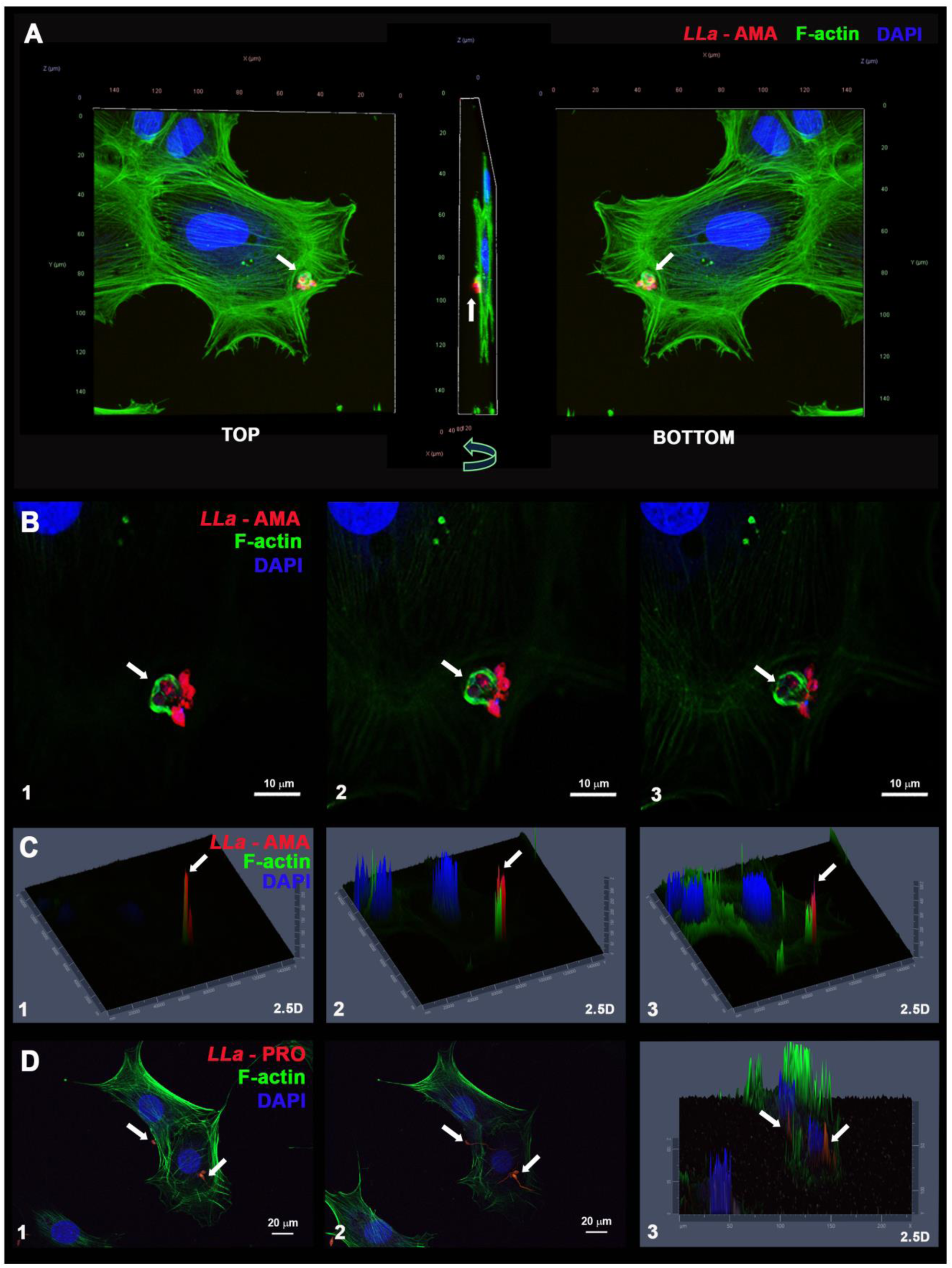
– *L. amazonensis* amastigotes induces localized actin polymerization at at invasion sites in MEFs. MEFs were incubated with *L. amazonensis* amastigotes (*LLa*-AMA, red) for 60 min, fixed with PFA and stained with phalloidin-A488 to visualize host cell F-actin (green) and DAPI to visualize the nuclei (blue). **(A)** Top and bottom views of a MEF during infection by *L. amazonensis* amastigotes (*LLa*-AMA, red) showing an actin ring (F-actin) surrounding the invading parasites. **(B)** Magnified views of the parasites indicated by the white arrow in A showing three different focal planes **(1 to 3)** of the site of invasion and the local actin rearrangement (white arrows). **(C)** 2.5D projection of figure B (1 to 3) showing amastigotes being internalized into MEF. **(D)** Same experiment but using *L. amazonensis* promastigotes (*LLa*-PRO, red). (**1**) Maximum intensity projection of MEFs during invasion by promastigotes; (**2**) A slide (single focus plane) showing that actin polymerization (F-actin) is not induced during promastigote internalization (white arrows). (**3**) 2.5D projection of image D. Images were taken with LSM 880 microscope/Airyscan detector (ZEISS).

**Fig. S5.**
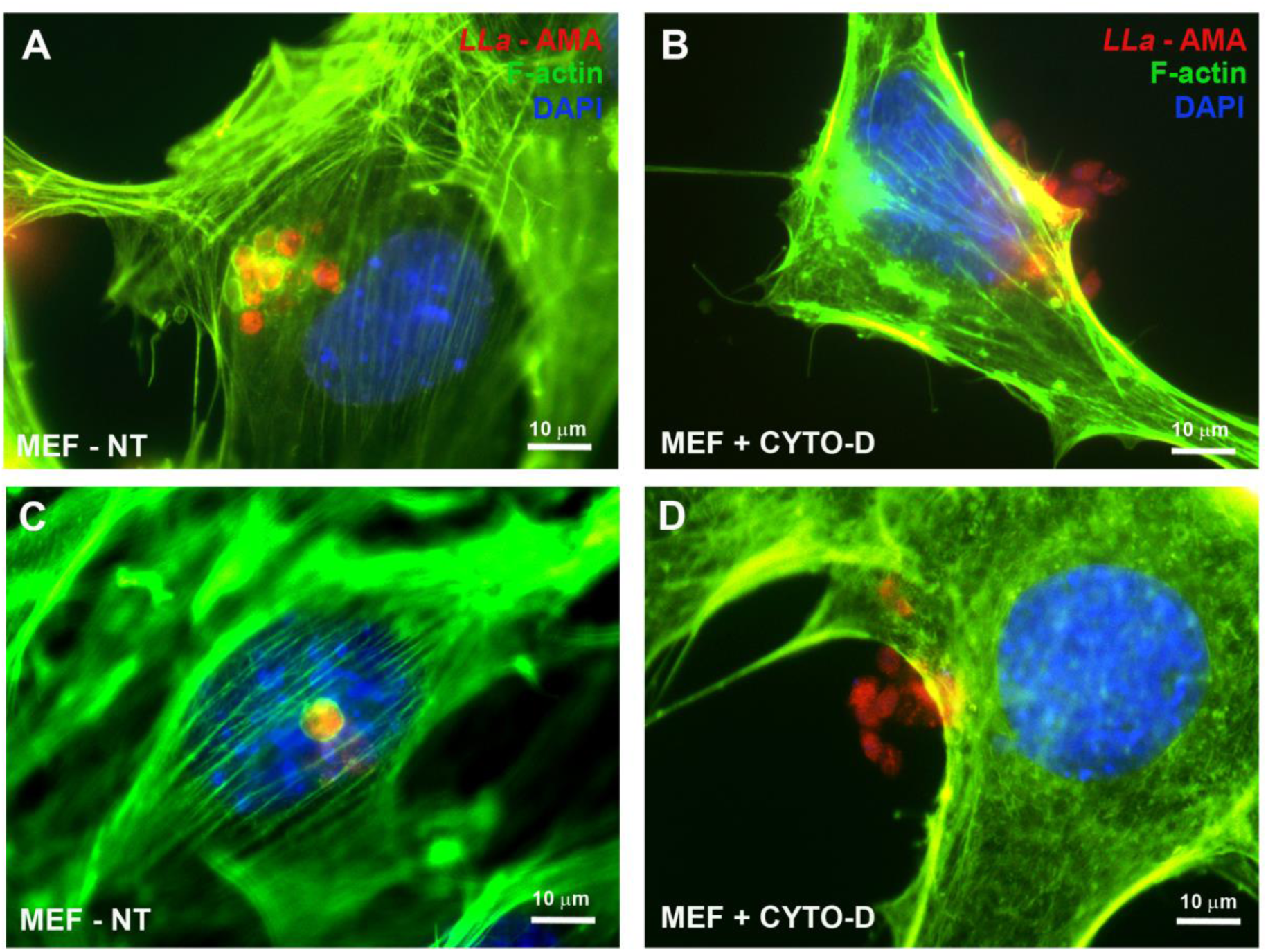
– Host cell actin drivers *L. amazonensis* amastigote invasion in MEFs. (Repeats of Fig. 6 D and F). Fibroblasts were infected with *L. amazonensis* RFP amastigotes (*LLa*-AMA, red) and analyzed by fluorescence microscopy. **(A and C)** Fibroblasts infected with LLA-RFP amastigotes (red) and **(B and D)** fibroblasts pre-treated with cytochalasin-D and infected with *LLa*-AMA (red). The cells were fixed with 4% PFA and stainedwith phalloidin-A488 (green) to label F-actin cytoskeleton and DAPI (blue) to label nuclei. In cells non-treated cells **(A and C)** parasites were observed internalized and surrounded by F-actin filaments, whereas in cells treated with cytochalasin-D **(B and D)** parasites were unable to perform cell invasion, remaining in the extracellular milieu, adhered to the external face of MEF’s plasma membrane. Images obtained using a BX60 upright compound fluorescence microscope (Olympus).

**Fig. S6.**
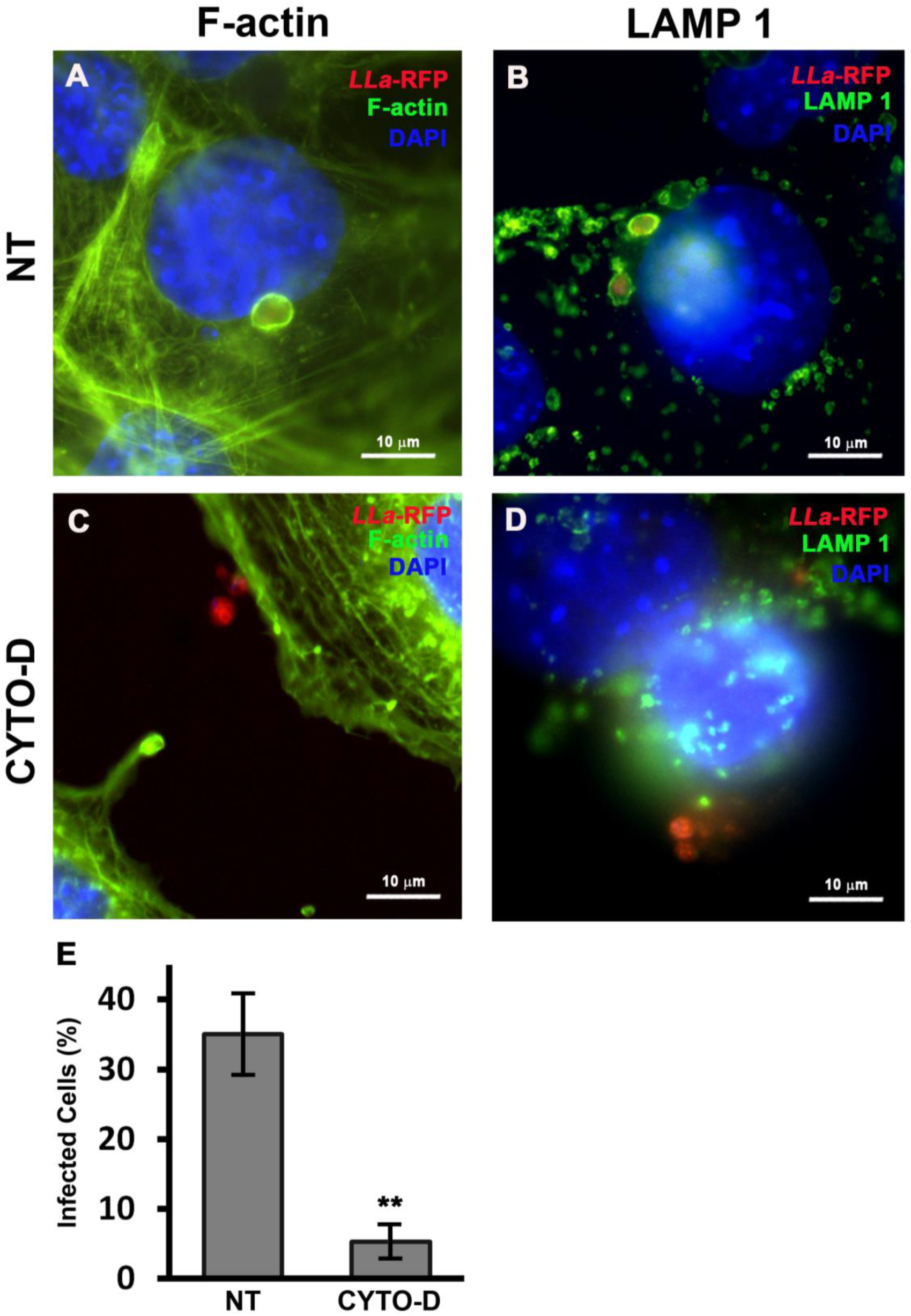
– Non-axenic *L. amazonensis* amastigotes are equally dependent on host cell F-actin cytoskeleton for invasion of MEFs. Macrophage-like cells (RAW cells) were infected with promastigotes and cultivated for 48 h. After parasite transformation into amastigotes and their replication within the host cell, the non-axenic amastigotes were obtained and incubated with (**A and B**) non-treated MEFs or (**C and D**) cytochalasin-D pre-treated MEFs. Cells were fixed with 4% PFA and stained with phalloidin-A488 (**A and C, green**) to label F-actin cytoskeleton or with **(B and D)** anti LAMP 1 antibody (green) to visualize lysosomes and parasitophorous vacuoles. **(E)** Infection quantification obtained by manual counting under the microscope. The data represent the mean±s.e.m of three independent biological replicates, * = 0,005, Student’s *t*-Test. Images obtained using a BX60 upright compound fluorescence microscope (Olympus).

**Fig. S7.**
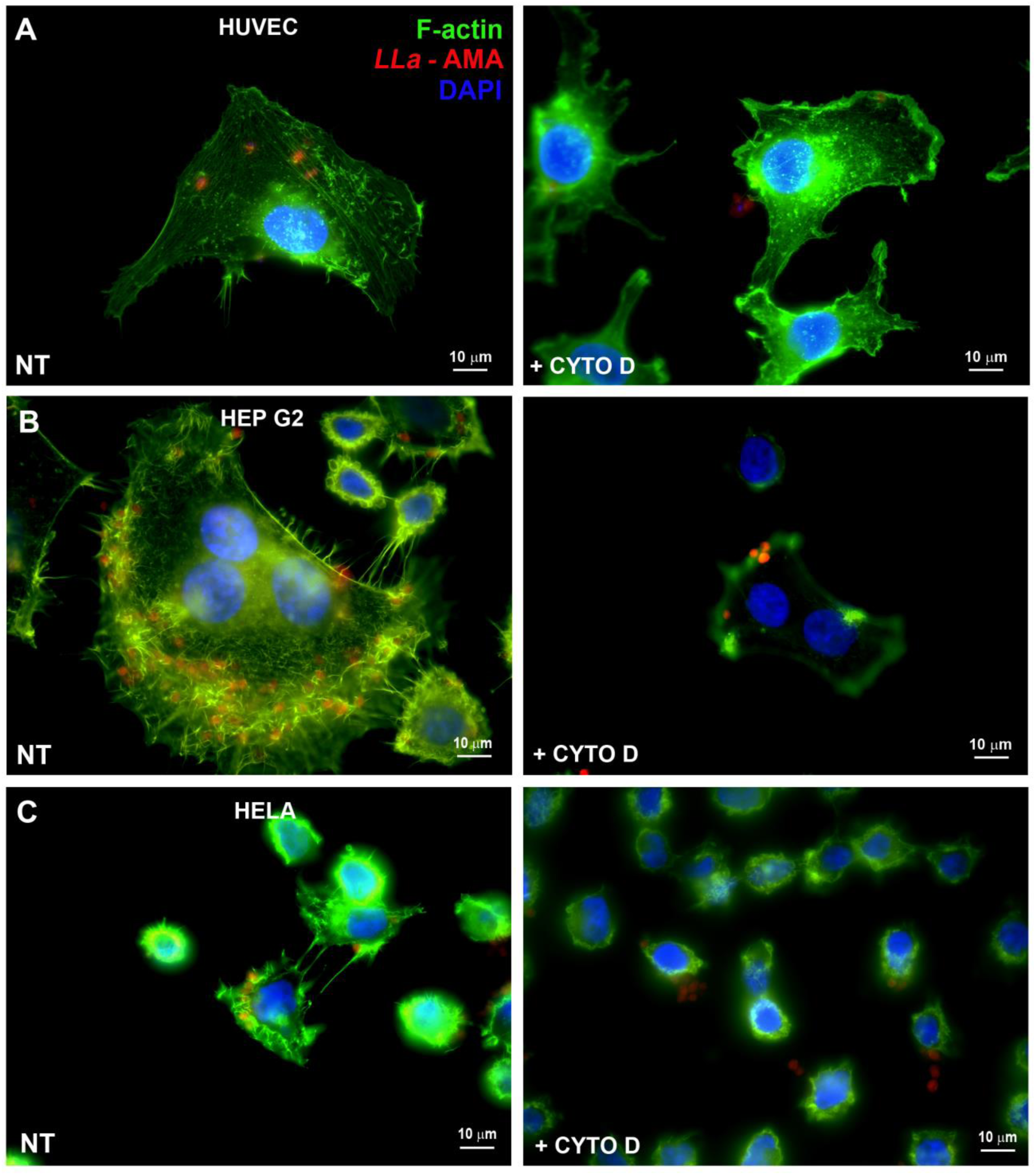
– Actin polymerization is required for host cell invasion by *L. amazonensis* amastigotes in different cell types. **(A)** HUVEC, **(B)** HepG2, and **(C)** HeLa cells were pre-treated with cytochalasin D (+CYTO-D) and then incubated with *L. amazonensis* amastigotes (*LLa*-AMA, red) for 120 minutes. Cells were subsequently fixed and stained with phalloidin-A488 (green) to label the F-actin cytoskeleton and DAPI (blue) to visualize nuclei. In non-treated control cells (NT), parasites were observed internalized and surrounded by F-actin filaments. In contrast, in cells pre-treated with cytochalasin D, amastigotes were unable to invade and remained attached to the cell surface. All images were acquired using a BX60 upright compound fluorescence microscope (Olympus).

**Fig. S8.**
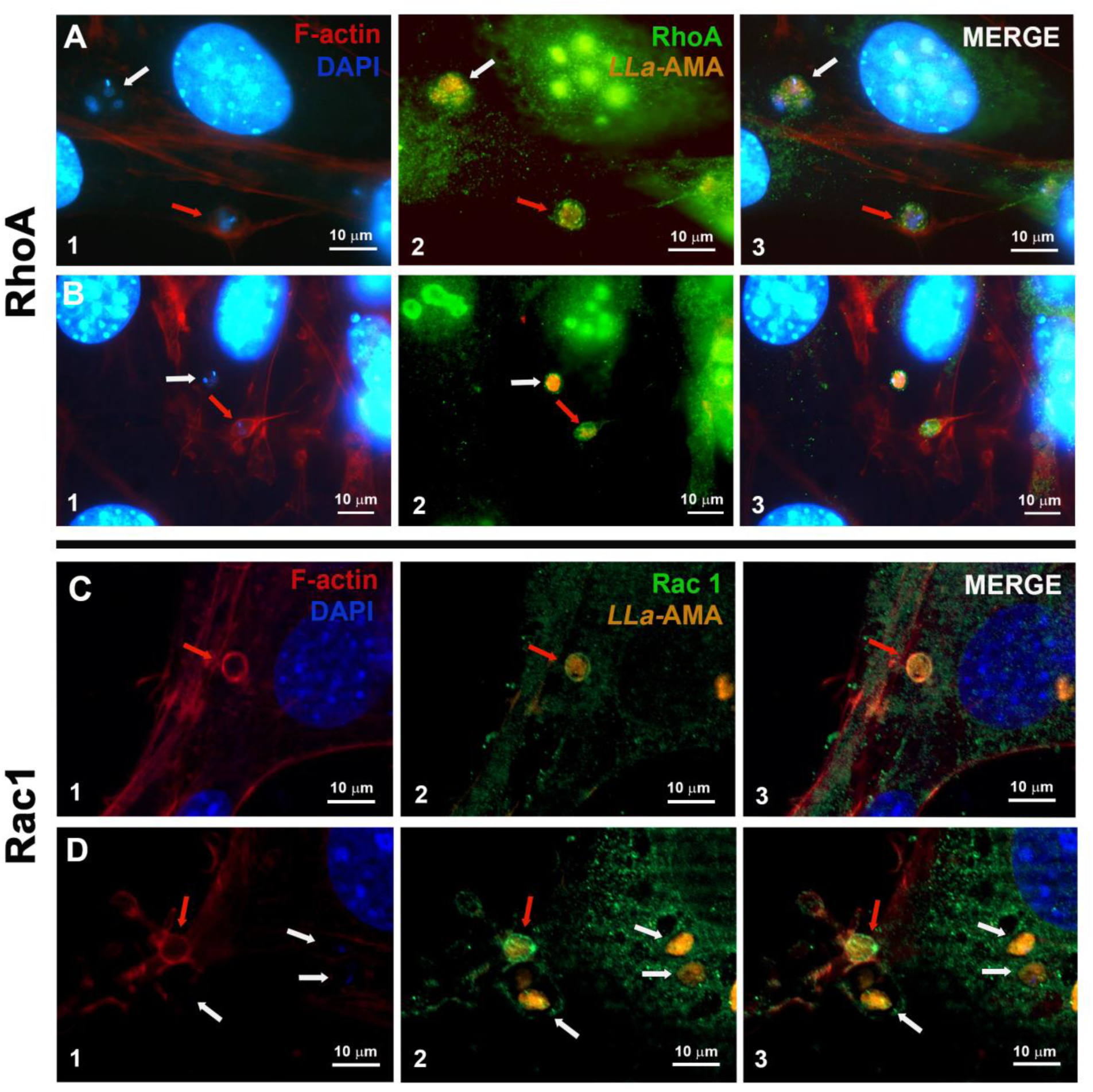
– The proteins of Rho GTPase family RhoA and RAC1 are recruited to *L. amazonensis* amastigotes invasion foci in MEFs. (**A-D**) MEFs were incubated with *L. amazonensis* amastigotes (*LLa*-AMA, orange) for 60 minutes. Cells were stained with phalloidin-Alexa Fluor 633 to visualize host cell F-actin (red), anti-RhoA or anti-Rac1 antibodies to detect the presence of these GTPases (green), and DAPI (blue) to label nuclei. (**A1**, **B1, C1 and D1)** show actin polymerization at sites of parasite entry (red arrows), which co-localizes with **(A2 and B2)** RhoA and **(C2 and D2)** Rac1. White arrows indicate parasites at later stages of infection, residing in vacuoles that lack F-actin accumulation and exhibit weaker labeling for both GTPases. **(A3, B3, C3 and D3))** show the merged images. Images were taken using confocal LSM 880 microscope (ZEISS). These are representative images of repetitions for the experiments shown in Fig. 7A-B.

**Fig. S9.**
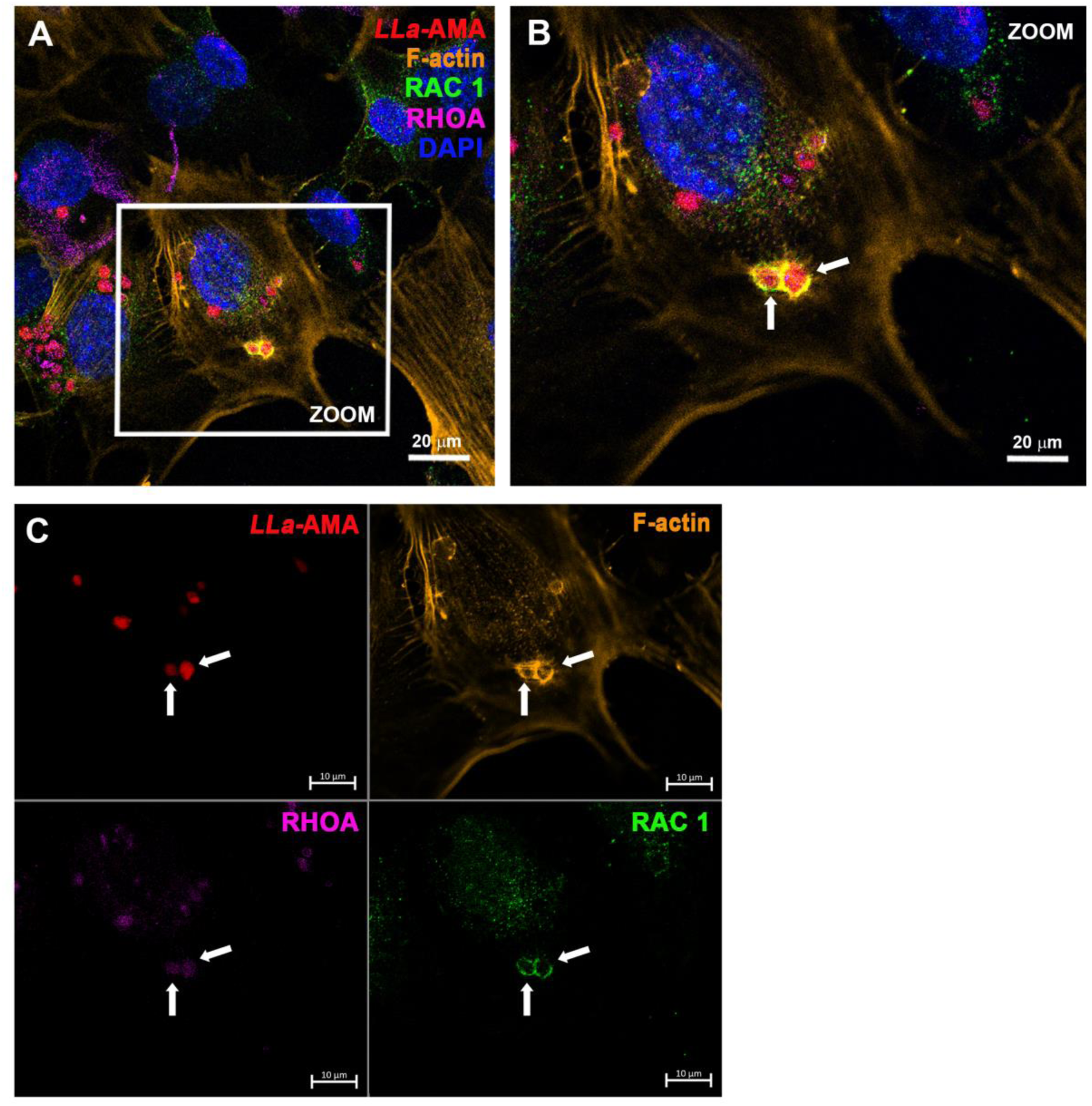
– RAC1 and RhoA are both present in F-actin-rich sites during the invasion of MEFs by *L. amazonensis* amastigotes. **(A)** Fluorescence microscopy analysis of MEFs infected for 60 minutes with *L. amazonensis* amastigotes (*LLa*-AMA, red), stained with anti-RAC1 (green) and anti-RhoA (magenta) antibodies to assess the recruitment of these proteins to sites of parasite entry. F-actin was labeled with phalloidin-A555 (orange), and nuclei were stained with DAPI (blue). **(B)** Magnified view of the region highlighted in panel A (zoom). **(C)** Individual fluorescence channels showing the labeling pattern. White arrows point to amastigotes during invasion process. This result is a repetition of the experiment shown in Fig. 7C-G. Images were acquired using the spectral detector ChS1 on a confocal LSM 880 microscope (ZEISS).

## Notes

### Competing Interest Statement

The authors have declared no competing interest.

